# Assessment of Nucleic Acid Structure Prediction in CASP16

**DOI:** 10.1101/2025.05.06.652459

**Authors:** Rachael C. Kretsch, Alissa M. Hummer, Shujun He, Rongqing Yuan, Jing Zhang, Thomas Karagianes, Qian Cong, Andriy Kryshtafovych, Rhiju Das

**Affiliations:** Biophysics Program, Stanford University School of Medicine, Stanford, CA, USA; Howard Hughes Medical Institute, Stanford University, Stanford, CA, USA; Department of Biochemistry, Stanford University School of Medicine, Stanford, CA, USA; Department of Chemical Engineering, Texas A&M University, TX, USA; Eugene McDermott Center for Human Growth and Development, University of Texas Southwestern Medical Center, Dallas, TX, USA; Department of Biophysics, University of Texas Southwestern Medical Center, Dallas, TX, USA; Lyda Hill Department of Bioinformatics, University of Texas Southwestern Medical Center, Dallas, TX, USA; Eterna Massive Open Laboratory, Stanford, CA, USA; Harold C. Simmons Comprehensive Cancer Center, University of Texas Southwestern Medical Center, Dallas, TX, USA; Genome Center, University of California, Davis, CA, USA

**Keywords:** CASP16, RNA structure, DNA structure, nucleic-acid protein complex structure, nucleic-acid ligand structure, structure prediction, deep learning, multiple sequence alignment

## Abstract

Consistently accurate 3D nucleic acid structure prediction would facilitate studies of the diverse RNA and DNA molecules underlying life. In CASP16, blind predictions for 42 targets canvassing a full array of nucleic acid functions, from dopamine binding by DNA to formation of elaborate RNA nanocages, were submitted by 65 groups from 46 different labs worldwide. In contrast to concurrent protein structure predictions, performance on nucleic acids was generally poor, with no predictions of previously unseen natural RNA structures achieving TM-scores above 0.8. Even though automated server performance has improved, all top-performing groups were human expert predictors: Vfold, GuangzhouRNA-human, and KiharaLab. Good performance on one template-free modeling target (OLE RNA) and accurate global secondary structure prediction suggested that structural information can be extracted from multiple sequence alignments. However, 3D accuracy generally appeared to depend on the availability of closely related 3D structure templates, and predictions still did not achieve consistent recovery of pseudoknots, singlet Watson-Crick-Franklin pairs, non-canonical pairs, or tertiary motifs like A-minor interactions. For the first time, blind predictions of nucleic acid interactions with small molecules, proteins, and other nucleic acids could be assessed in CASP16. As with nucleic acid monomers, prediction accuracy for nucleic acid complexes was generally poor unless 3D templates were available. Accounting for template availability, there has not been a notable increase in nucleic acid modeling accuracy between previous blind challenges and CASP16.

## Introduction

Nucleic acid (NA) structure prediction has lagged behind the transformative advances in protein structure prediction, as measured in the RNA-Puzzles and CASP blind competitions^1–6^. Two years ago, CASP15 revealed the underperformance of deep-learning based algorithms for RNA compared to human experts^6^. Since CASP15^7^, the RNA structure community standardized databases^8,9^; improved self-assessment through more stringent train-test splits^10^; integrated deep-learning-based methods with physics-based methods, evolutionary information^11,12^, and functional data; and increased the amount of training data by accelerating experimental structural determination^13,14^, self-distillation^15^, and high throughput, low-resolution experiments^16–19^. In addition, the AlphaFold 3 server, which extends AlphaFold modeling to nucleic acids and their complexes, was released^20^.

These events, along with the prospect of atomic-accuracy nucleic acid structure prediction, have led to increased participation from both experimental structure providers and predictors in CASP16 compared to CASP15. As a result, CASP16 presented the opportunity to rigorously benchmark nucleic acid prediction across difficulty levels, across prediction methodologies, and even across molecular types, with targets involving DNA structure as well as complexes involving RNA-ligand, RNA-protein, DNA-protein, and RNA-RNA interactions. Here, we present an assessment of blind predictions across all NA targets of CASP16. Complementary to this assessment, other analyses have evaluated subsets of these targets with particular attention to functionally relevant regions assigned by structure providers^21^, protein domains of complexes^22^, water and ion networks^23^, conformational ensembles^24^, and classic RNA-specific features developed in RNA-puzzles^25^.

Our assessment across all targets has revealed that NA structure prediction accuracy remains dependent on the availability of previously solved experimental 3D structural templates, with little to no success when a similar structure is unavailable. Additionally, comparison across the history of RNA structure prediction indicates that the recent increase in interest in RNA structure prediction and integration of deep learning has yet to result in a significant increase in prediction accuracy. Nevertheless, there is cause for optimism. For example, the global fold of a completely novel RNA was predicted correctly, although details of tertiary and quaternary contacts were not correctly captured. Perhaps most promisingly, automated servers were able to improve on the best available structure templates in some cases, although both servers and human experts need to make major progress before achieving accuracies comparable to experimental precision.

## 2 | Methods

### 2.1 | Target Selection

In CASP16, NA targets were assessed in four categories: NA monomers, RNA-only multimers, hybrid NA-protein complexes, and NA-ligand complexes. Each prediction target was assigned a target label; RNA structure labels started with “R1”, DNA structure labels started with “D1”, and hybrid target labels started with “M1”.

With the introduction of multimeric NA-containing complexes as targets, whether the RNA or DNA chain alone should be assessed as a NA monomer target was decided on a case-by-case basis. We excluded targets where the NA structure was simple or heavily scaffolded by proteins: simple double helical structures (M1216, M1228, M1239, M1268, M1282, M1287), a structure with a template and extensive scaffolding by proteins (M1297, a SPARSA gRNA-DNA complex), and a target where the NA was small (< 10 nt) and scaffolded by a protein (M1276, a cryptic DNA-binding protein UDE). We additionally split two RNase P-tRNA complexes, each with a protein and two RNA chains of different sequences into two NA assessment units (R1221s2 and R1221s3; R1224s2 and R1224s3). Finally, because all the RNA-only multimer targets were symmetric, we only evaluated one chain in the monomeric assessment. The two conformations of R1253 (ROOL RNA) were scored as separate targets. The three stoichiometric states of R1283 (*Enterococcus* ROOL RNA), had very similar monomeric structures, so only R1283v1 was assessed for monomer prediction tasks. This resulted in 36 targets in the NA monomer category. A DNA target D1273 (dopamine aptamer) was not considered for the RNA base-pair and motif analysis, multiple sequence alignment (MSA) analysis, or template analysis, resulting in 35 targets for those analyses.

All targets containing a small molecule ligand bound to a NA were considered for the NA-ligand assessment. Out of six targets of this type, one (R1262) was released with the incorrect bond order and thus was not assessed. Furthermore, two targets (R1288 and D1273) were not considered for the NA-ligand ranking because NA-ligand interactions were difficult to evaluate in the context of poor NA-pocket predictions. Hence, ranking was based on only three targets: R1261, R1263, and R1264. Ions and water were not included in this NA-ligand assessment, but are considered in a separate paper submitted to this CASP16 special issue^23^.

All NA multimer targets were considered in the RNA multimer category, resulting in 11 targets. Additionally, all targets containing NA and proteins were selected for the hybrid category, resulting in 18 targets. For RNA multimers and hybrids, some targets were also considered for a “Round 0” prediction where predictors were not told the stoichiometry of the complex (**Table 1**). These Round 0 predictions were labeled “R0” and their prediction time of two weeks was immediately followed by the Round 1 prediction, “R1”, where predictors were now told the correct stoichiometry and could submit predictions again.

**Table 1:**
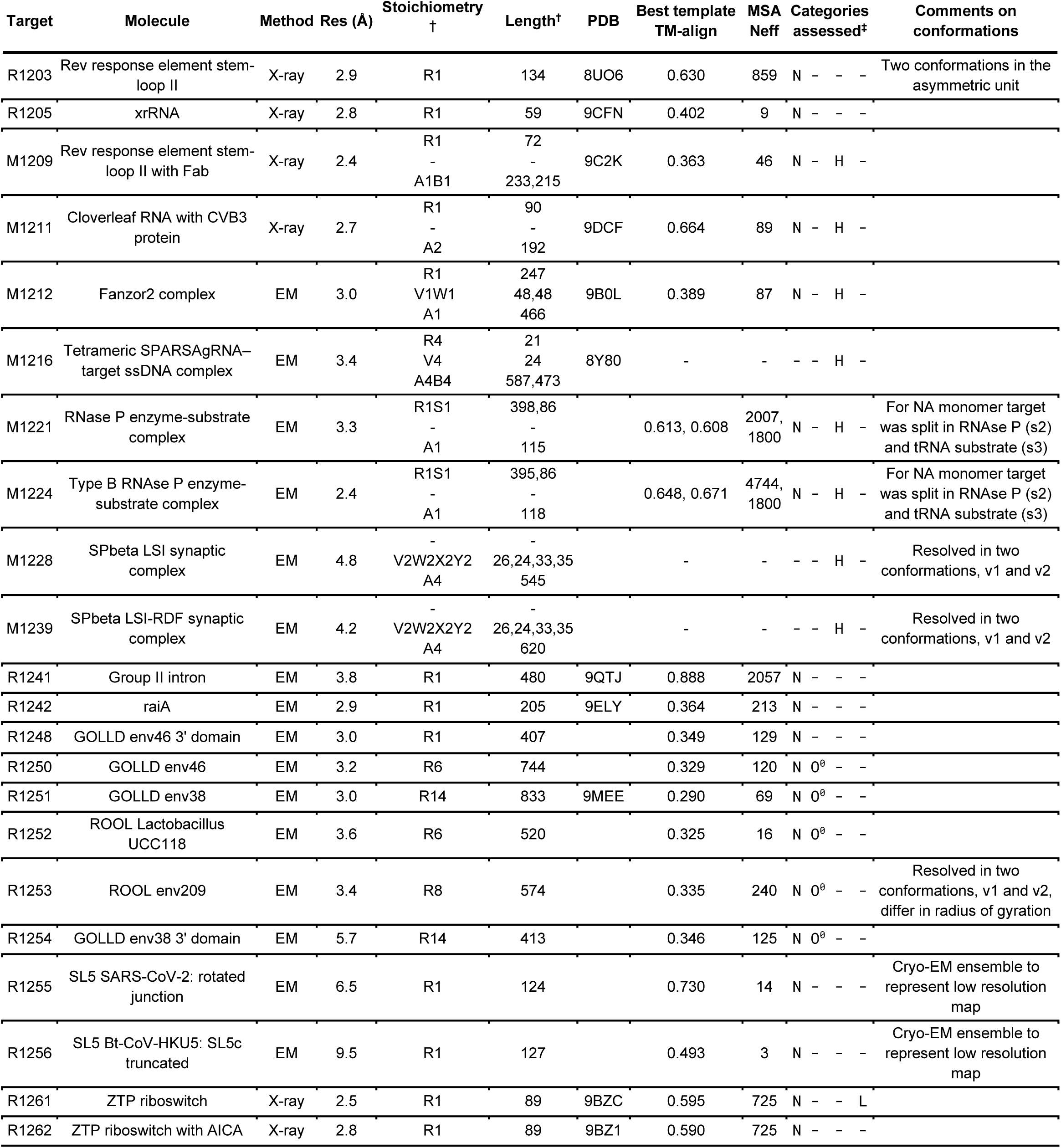
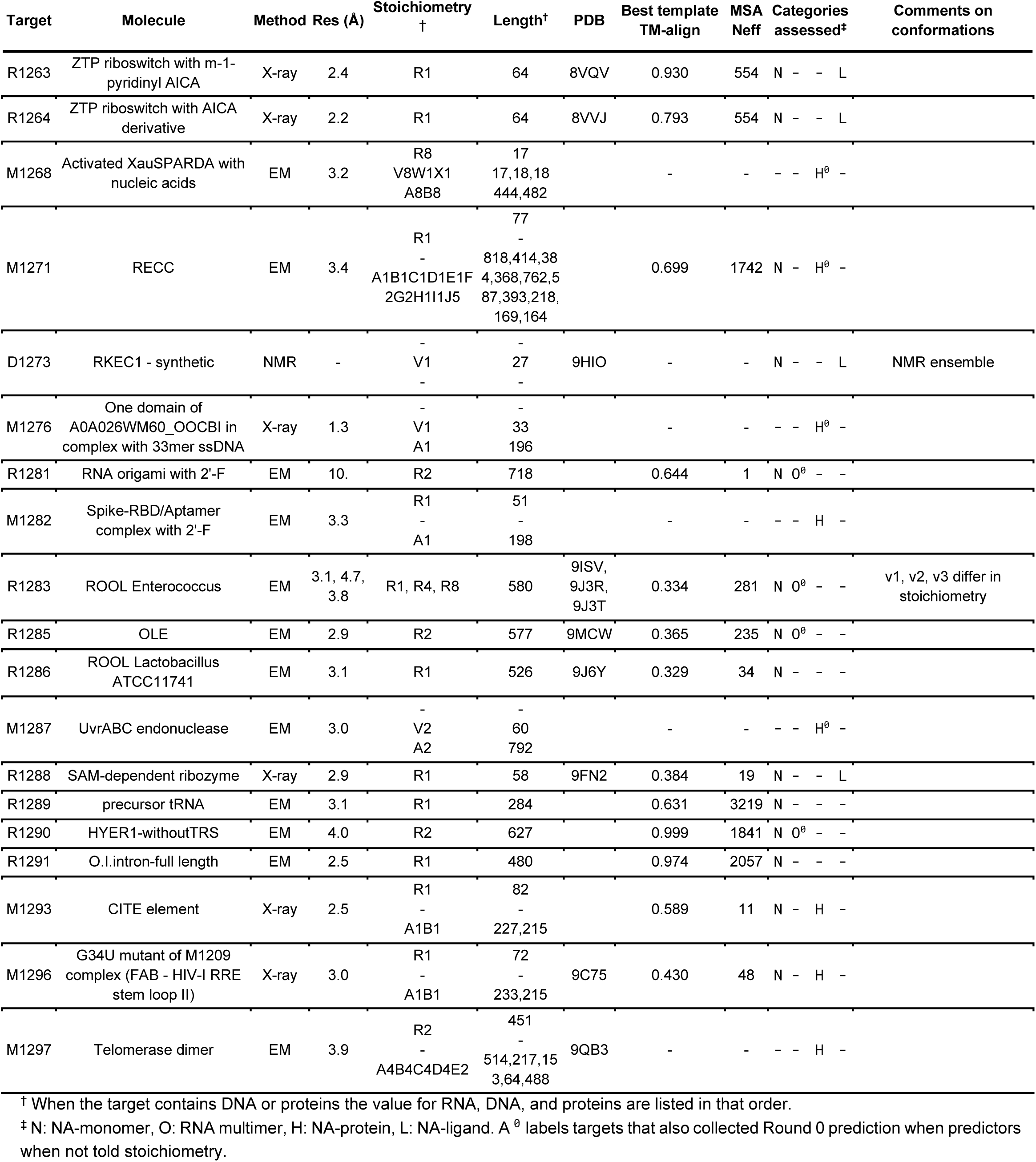
Summary of NA-containing targets.

### 2.2 | Identification of Best Template

The best structure template for each blind prediction target in CASP16, as well as in prior blind trials RNA-Puzzles and CASP15, was identified using US-align (Version 20220511) with the command line option “-mol RNA” and the wrapper script USalign.py (available at https://github.com/daslab/daslab_tools/). This template search was based on structural similarity and does not take sequence similarity into account. To allow for the direct comparison of scores for each reference structure against the best previously available template and against the best blind prediction, US-align was run with the default sequence-independent alignment option (referred to here as ‘TM-align’ score), since potential templates have different sequences than the target sequence. For each reference structure, the TM-align scores amongst PDB depositions whose release dates predated the target deadlines were computed. The best template was then selected by TM-align score. As potential templates, all depositions in the PDB containing at least one RNA chain released prior to the date of November 11, 2024, were identified using the PDB search tool (https://www.rcsb.org/search) and downloaded as mmCIF formatted files with the PDB-provided batch_download.sh script (https://www.rcsb.org/scripts/batch_download.sh). Release dates were extracted using get_cif_info.py (available at https://github.com/daslab/daslab_tools/) which looks at the ‘revision_date’ tags in the mmCIF file. The reference structures for CASP15 and CASP16 were the ones made available to assessors at the time of their respective assessment; structures for the RNA-Puzzles were downloaded from rnapuzzles.github.io. Information on the target deadlines and server status were compiled from the CASP15, CASP16, and RNA-Puzzles websites.

### 2.3 | Measurement of Multiple Sequence Alignment Depth

The normalized number of effective sequences (Neff) quantifies the number of non-redundant sequences, here defined as sequence identity less than 80 %, in an MSA. For each RNA monomer target, the full monomer target sequence was used to calculate an MSA using rMSA^26^ with the RNAcentral^27^ (v20.0, 2022-03-28), RFam^28^ (v14.7, 2021-12-09), and NCBI Nucleotide (2022-10-03) databases. The sequence identity between each pair of sequences in an MSA was calculated after ignoring any positions that were gapped in both of the sequences. For each sequence in an MSA, the weight for that sequence was calculated as the number of other sequences in the MSA with a sequence identity above 80 %. This threshold was previously used by AlphaFold2^29^ and trRosettaRNA^30^. The Neff value was then calculated as the sum of inverse weights for each sequence in the MSAs. The results were compared to different implementations of Neff, Neffy^31^: the symmetric method of Neffy differs because it counts matched gaps and the non-symmetric Neffy method because it is not symmetric across pairs of sequences. While numerical differences arose, the trends observed were the same. Additionally, using a 62 % sequence identity threshold, as used in the proteins assessment^32,33^, reduces all Neff values, but does not affect the trends.

### 2.4 | Computation of Full Structure Metrics

All predictions were scored against the provided experimental structures. When multiple experimental structures were available, the predicted model was compared to all experimental structures and was assigned the best score for each metric across all experimental structures.

The following programs were used to calculate metrics: (1) US-align^34,35^ for TM-score (with sequence correspondence) and TM-align (without sequence correspondence); (2) Local-Global Alignment^36^ for GDT_TS with sequence correspondence, where C4’ atoms were used for RNA and DNA; (3) OpenStructure^37^ for local distance difference test (lDDT)^38^ with sequence correspondence and with steric penalty; (4) rna-tools^39^ which uses ClaRNA^40^ for Interaction Network Fidelity (INF) scores.

### 2.5 | Group Labeling

Groups in the nucleic acid monomer prediction category were labeled according to prediction strategies described by the groups. Server and human groups are differentiated in the amount of time the group had for prediction: 3 days for servers, 2 weeks for humans. All other information about groups was obtained through abstracts, presentations at the CASP16 conference, and, when applicable, manuscripts the group cited in their abstract and/or released after the CASP16 competition describing their strategies. While many groups did self-label (e.g., marked MSA as Y or N), this labeling was not always clearly applicable to their RNA modeling strategies because the same abstract was used to describe protein structure prediction and/or multiple methods from the laboratory. Instead, we manually labeled traits of the methodology, marking traits as N/A when the abstract was absent or did not contain information regarding that category. Many groups used multiple methods to, for example, generate models. In this case, the group was marked as using a technique if any of the methods they used employed that technique. A group was marked as using MSAs if they created an MSA, manually looked at an MSA, or their method implicitly used MSAs. A group was marked as using templates if they or an algorithm identified PDB templates; simple, small fragments were not counted as templates.

The AlphaFold 3 server^20^ is used as a baseline prediction throughout. AlphaFold3 predictions were submitted by Arne Elofsson as group TS304. Each target sequence was submitted to the server and all 5 output models were uploaded as the prediction. The AlphaFold3 server participated blindly: assessors were not aware of the participation of the AlphaFold3 server nor were they aware of the group number until after assessment was completed.

### 2.6 | Computation of Interface Metrics

OpenStructure was used to calculate all interface metrics for multimeric complexes. For NA-NA, protein-protein, and NA-protein interfaces the F1-score (ICS) and Jaccard coefficient (IPS) of the inter-chain pairs of residues^41,42^, the lDDT of interface residues (i-lDDT), and the DockQ^43^ were calculated. For hybrid targets, each interface was labeled as NA-NA, NA-protein, or protein-protein. To calculate the NA-containing interface scores, the per-interface NA-NA and NA-protein scores were summed, weighted by the number of residues in the interface. These NA-containing interface scores were used for ranking. The following models were excluded from analysis: R1254 from group 033, because all atoms after the first chain were placed at the origin (**Supplemental Table 1**).

For NA-ligand interfaces, the i-lDDT of the NA-ligand interface (labeled as LDDT-PLI for protein ligand interface, labeled here simply as i-lDDT), lDDT of the ligand-pocket nucleotides (lDDT pocket), the Root-Mean-Square-Deviation (RMSD) of the ligand-pocket nucleotides, and the Binding-site superposed Symmetry-corrected pose Root-Mean-Square-Deviation (BiSyRMSD) of the ligand pose were calculated using OpenStructure^44–46^. The ligand pocket for R1263 and R1264 is formed from two symmetry pairs of the RNA, but predictors only predicted one chain. Hence, predictors could only predict partial binding sites with some predictors placing it in chain A of the target pocket and others in chain B of the target pocket. To account for this, the ligand metrics were calculated against the dimer ligand pocket interaction and then these scores were normalized by the score for the target half-pocket (chain A or chain B) against the target dimer.

### 2.7 | Ranking

For each CASP NA category, we used a ranking based on Z-scores, the number of standard deviations by which the model’s accuracy differed from the mean. In rankings we excluded the group 338 predictions for R1242 as they communicated that they had experimental data on the target. We also excluded groups that participated in less than 60% of the targets in the category. In some cases, the experimental reference structure may not be the only structure the NA adopts in solution. To account for this and to add leniency for the currently modest levels of accuracy in NA structure prediction, all five submitted models were considered. For each metric, the best score amongst all five models was considered. The mean and standard deviation of scores for each metric were calculated based on the top score from every group. To avoid very poor predictions skewing Z-scores, scores 2 or more standard deviations below the initially calculated mean were considered outliers, and the mean and standard deviation were recalculated without these outliers^6^. The final Z-scores for ranking were then calculated for each of the four categories with the following weighting for different metrics:

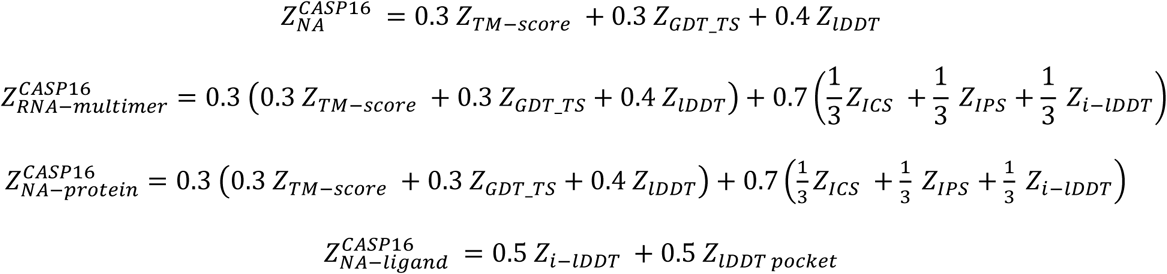

The Z-scores were summed across all targets in the category. To reduce over-rewarding for similar targets, we only considered the best score for targets with the same sequence, grouping the following sets of targets: [R1221s3 and R1224s3], [R1241 and R1291], [R1253v1 and R1253v2], [R1261 and R1262], [R1263 and R1264], [R1283v1, R1283v2, and R1283v3], [R1253v1o and R1253v2o], [R1283v2o and R1283v3o], [M1228v1 and M1228v2] and [M1239v1 and M1239v2]. The prediction of multiple structures, or the conformational ensemble of structures, is assessed elsewhere in this issue^24^. For all categories, poor predictions were not penalized: if a Z-score was below zero for a target, Z-score was set to 0 for summed ranking. All Z-scores were then summed across all categories to obtain a NA ranking.

Confidence intervals for each score were calculated based on 1,000 bootstrap replicates, selecting targets with replacement. Based on the bootstrapped scores, a 68.2 % confidence interval was displayed on plots. Confidence intervals were used instead of the standard error of the mean because the distributions were not normally distributed.

### 2.8 | Secondary Structure Analysis

Secondary structures were extracted from CASP16 models with DSSR (v1.9.9-2020feb06)^47^. Some models, in particular due to large clashes, failed to run (**Supplemental Table 1**). The base-pair list was extracted from the table in the output file directly because the dot-bracket structure produced by DSSR, in particular for multimers, can contain errors. The canonical base pairs were defined as those labeled as Watson-Crick-Franklin (WC) and wobble base pairs (hereafter referred to as ‘base pairs’ or ‘pairs’). All other base pairs are defined as non-canonical base pairs and analyzed separately. Crossed base pairs (pseudoknots) were defined as non-nested canonical base pairs, that is any canonical base pair (*i*,*j*) for which another canonical base pair (*k*,*l*) exists with *i* < *k* < *j* < *l* or *k* < *i* < *j* < *l*. Singlet base pairs were defined as any canonical base pair that was not part of a stem, that is (*i*,*j*) such that there was no neighboring canonical base pair between *i* +*1* and *j* – *1* or between *i* – 1 and *j* + 1. Intermolecular base pairs were identified as any canonical base pair between nucleotides in different chains.

The base pairs, crossed pairs, singlet pairs, and non-canonical pairs were scored using F1-scores (see below). The predicted pair was considered a match to a target pair in the experimental reference structure if it involved exactly the same two nucleotides. In the future, non-canonical base-pair prediction could additionally be evaluated on the ability to predict the correct base-pair edges, though this was not explored here due to inconsistencies in edge assignments by different software tools in some cases. Any predicted base pair containing at least one nucleotide that was unresolved in the experimental reference structure was ignored, that is it did not count for or against the precision of the prediction.

For each model, for each interaction type, the precision is the number of interactions predicted correctly divided by the total number of interactions predicted, the recall is the number of interactions predicted correctly divided by the total number of interactions in the target, and the F1-score is the harmonic mean of the precision and recall. If the target reference structure and prediction had no instances of an interaction type, then the model received no score and hence did not contribute to the ranking. To penalize for overprediction, if the reference structure had no instance of that interaction type but the prediction did, a value of 0 for the F1-score was recorded. Additionally, if participants did not submit a model, they were assigned an F1-score of 0. Like the Z-score ranking, the base-pair F1-scores were considered for all 5 submitted models and the top score for each group was selected. When a target had multiple references, each prediction was compared to all references and the best score was taken; a null score, or correctly predicting no interaction, was considered the best score. For each group, the top F1-score for an interaction type was averaged over all targets, ignoring null scores.

### 2.9 | Generation of Benchmark Secondary Structure Predictions

There are numerous packages for predicting RNA secondary structure, including pseudoknots, from sequence alone, without explicit modeling of 3D coordinates or requiring MSAs. Using the OpenKnot Score Pipeline (https://github.com/eternagame/OpenKnotScorePipeline), we ran ViennaRNA (2.4.16)^48^, CONTRAfold (v2, last commit June 19, 2017)^49^, EternaFold (last edit October 28, 2021)^50^, RNAstructure Fold program (v6.4)^51^, HotKnots (v2.0)^52^, IPknot (last commit June 9 2021)^53^, SPOT-RNA (last commit April 1 2021)^54^, RNet^17^ (three variants: (1) the RNet release model, trained on the experimental SHAPE and DMS data from the Ribonanza dataset and, as a form of distillation^55^, average predictions of the 3 top models entered in the Ribonanza Kaggle competition; (2) RNet-All, which was further fine-tuned on experimental SHAPE and DMS data for the Ribonanza test set; and (3) RNet-HQselect, which was trained solely on experimental SHAPE and DMS data for the Ribonanza training sequences, revisited with higher sequencing depth), and Shapify-HFold (last commit February 15, 2022)^56^. Knotty (last commit March 28 2018)^57^, NUPACK (3.0.6)^58,59^ and pknots (last commit January 30, 2020)^60^ were all run with the pseudoknot option on. However, these algorithms were too memory-or time-intensive for long sequences, so NUPACK was run without pseudoknots for sequences longer than 240 nucleotides; pknots was run without pseudoknots for sequences longer than 250 nucleotides; and pknots predictions were substituted in for Knotty, which cannot be run without pseudoknots for sequences longer than 450 nucleotides. Further, we generated base pair probability matrices using ViennaRNA, EternaFold, and CONTRAfold, and used a linear-sum assignment algorithm^61^ or a ThreshKnot heuristic^62^ to generate potentially pseudoknotted structures. None of these algorithms use MSAs, although some have been trained on known secondary structures or many sequences.

### 2.10 | Identification of Structural Motifs

Rosetta rna_motif^63^ was used to identify tertiary RNA motifs in PDB structures. The RNA motifs analyzed here were: A-Minor, GA-Minor, T-Loop, Intercalated T-Loop, Loop-E Submotif, GNRA Tetraloop, Tetraloop Receptor, Platform, Tandem GA Sheared, UA handle, Bulged G, U Turn, and Z Turn. Additionally, the docked A in A-Minor interactions and the intercalated nucleotide in the intercalated T-loop (also called T-loop/D-loop or T-loop/T-loop interactions) result in long-range tertiary contacts that are of particular interest in 3D structure prediction; these two kinds of interactions were separately evaluated as an additional prediction test. Tandem GA Watson-Crick interactions and double T-loops were identified by rna_motif, but none were found in any target or prediction and hence were not analyzed. The PDB reference structures for three targets were amended for rna_motif to run: segids were removed from R1264 and R1296, and two residues (29 and 371) were removed from R1285. 182 model structures could not be processed with Rosetta rna_motif and thus were excluded from this analysis (**Supplemental Table 1**). Intermolecular motifs were identified as any motifs involving nucleotides from more than one chain. A motif was deemed to be predicted correctly if it contained the same nucleotides as the target motif in the experimental reference structure, and each nucleotide occupied the same chemical positions as defined by rna_motif. This approximation was appropriate for the dimeric and symmetric homo-oligomeric complexes in CASP16, but in the future, for more complex targets and more accurate predictions, the chain mapping problems should be considered^64^. The choice of best score and calculation of F1 scores for motifs were computed as with the F1 base pair metrics described above.

### 2.11 | Assessment of Stoichiometry (Round 0) and Symmetry (Round 1)

For Round 0 predictions of RNA multimers and hybrids, we reported the percentage of predicted stoichiometries that were the same as the experimentally derived stoichiometry. When there were multiple experimentally derived stoichiometries for a target, all were considered correct.

For RNA multimer targets, we additionally assessed symmetry predictions. Potential symmetries for each RNA multimer model were collected from AnAnaS^65^. To run RNA PDBs in AnAnaS, each RNA nucleotide was relabeled as an alanine and the phosphorus atom was relabeled as C-alpha. From all potential symmetries with a 10 Å RMSD cutoff, the highest order symmetry was determined using two criteria: the most chains involved and the most symmetry elements (i.e., a D4 dihedral symmetry would be selected over a C8 cyclic symmetry). As with stoichiometry, we reported the percentage of predicted symmetries that were the same as the experimentally derived stoichiometry.

### 2.12 | Identification of Independently Determined Structure Pairs

To identify the best possible TM-align values that might be achieved given RNA molecules’ inherent flexibilities and experimental uncertainties, a manually curated set of structure pairs was collected. The following criteria were used: (1) the structures in each pair were solved by independent experimental groups, (2) the structures did not represent distinct conformational states of the RNA as determined by expert knowledge, (3) the structures had the same sequence, allowing for minor mutations used during the experimental process (e.g., mutations made to enable crystallization or observation by cryogenic electron microscopy).

## Results

### 3.1 | Expansive Range of Nucleic Acid Containing Targets

CASP16 challenged 65 predictors with a total of 42 NA containing targets, provided by 22 experimental groups^21^. For one target, R1260, CASP asked groups to predict water and ion ensembles, a distinct and new assessment that is discussed elsewhere in this issue^23^. We assessed 4 NA categories: 1 DNA and 35 RNA targets as nucleic acid monomers, 11 RNA-only multimers, 18 hybrid complexes (complexes containing protein and DNA and/or RNA), and 5 RNA ligand interactions (**Table 1** and **Figure 1**). Together, these four categories informed a broad understanding of prediction accuracy of NA containing biomolecular systems.

**Figure 1:**
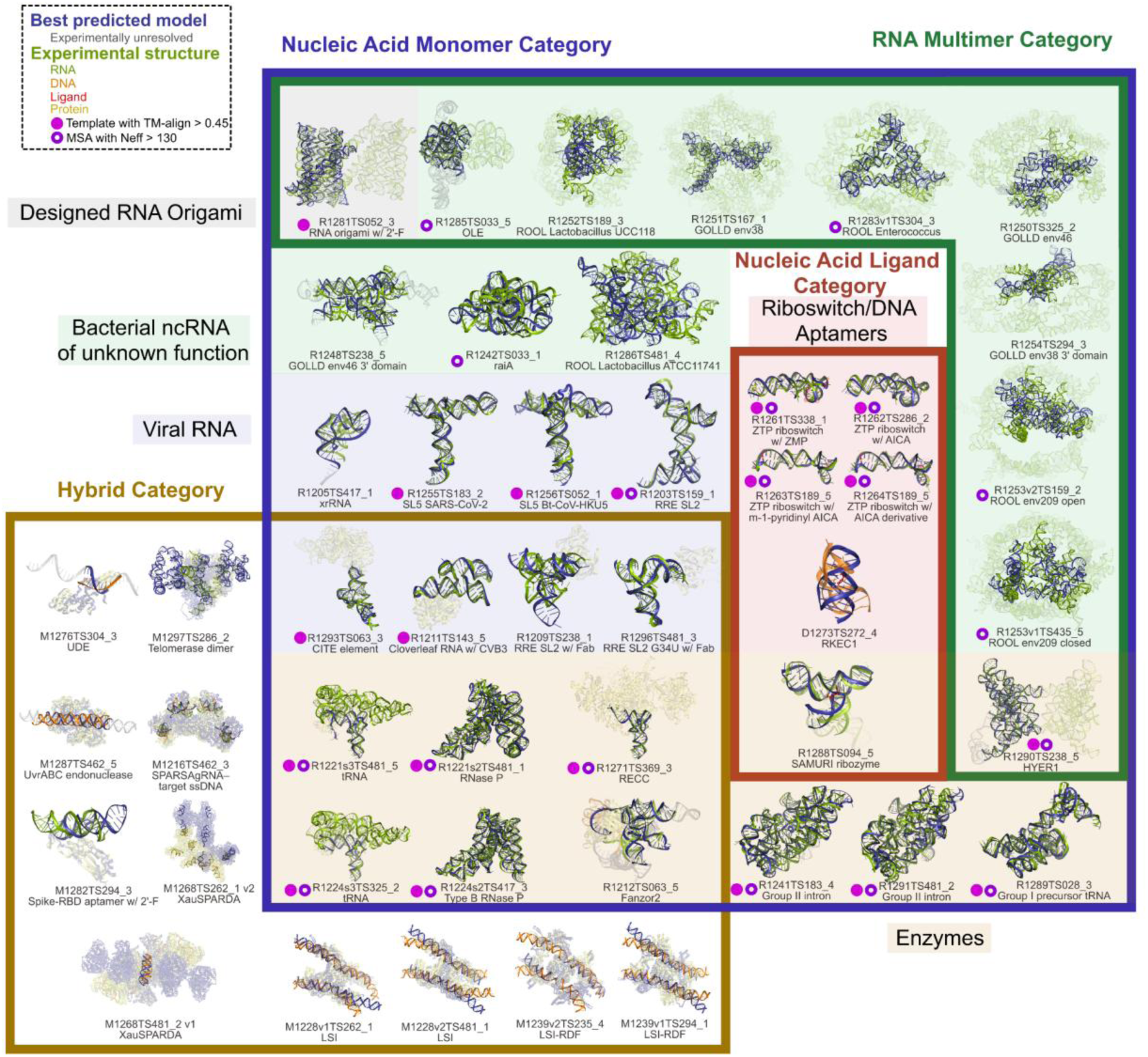
Nucleic acid targets assessed in CASP16. The RNA in each target is displayed in green, the DNA in orange, the ligand in red and the protein in yellow. The best predicted model by TM-score is overlaid and displayed in dark blue. Residues that were predicted but not experimentally resolved are colored gray. For RNA multimers only one predicted chain is shown and other chains in the target are made transparent. Likewise the protein chains are made transparent. The best predicted model is labeled under the image. The targets are grouped by the categories in which the target was assessed (boxed regions). Targets in the NA monomer category are additionally grouped by functional role (shaded regions), and if a target has a template (TM-align > 0.45) it is labeled with a pink filled dot. If a target has a deep MSA (Neff > 130) it is labeled with a purple open dot.

An important goal of CASP is to benchmark which structure prediction problems are easy and which are difficult for the field. In order to survey this, we decided to include a full range of targets from “easy” targets that tested their ability to refine structures from templates to “hard” targets with no templates that tested groups’ ability to predict structure from sequence, evolutionary information, and functional information alone. Additionally, the easy targets served as controls for assessment metrics, particularly in new categories like RNA multimer assessment.

### 3.2 | Metrics to Assess NA Monomer Structure Prediction

The expansion of the number and type of NA-containing targets motivated the identification of metrics that were independent of biomolecular type and hence could enable a comparison of performance across RNA, DNA, and protein targets. Previously, CASP15 scoring schemes used GDT_TS^36^ and TM-score^34,35^ to assess the accuracy of predictions’ global fold. These metrics can be calculated for all types of biomolecules. We noted that, in particular for large complexes, GDT_TS and TM-score can have different resolving capacities (**Supplemental Figure 1A**), so we decided to include both metrics for CASP16 rankings. For clarity, we have chosen to mainly report TM-score as it qualitatively matched our manual accuracy assessments and is available in a sequence-independent version (here called TM-align) that allows for comparison to templates with different sequences.

Compared to global fold metrics, metrics that measure the local accuracy can be important for differentiating predictions with highly inaccurate or highly accurate global folds. For the prediction of local accuracy, the lDDT^38^, INF (here the INF score refers to the INF_all score)^66^, and clashscore^67^ were used in CASP15. The INF score is a metric that identifies base-pairing and stacking interactions. While this metric could be developed further for the assessment of DNA, DNA-DNA, DNA-RNA, and RNA-RNA interfaces, it would not be suited for RNA-protein interfaces. On the other hand, the lDDT and clashscore are applicable to any molecule. There is a strong correlation between INF and lDDT (**Supplemental Figure 1B-C**); while not specifically evaluating stacking and hydrogen bonds, lDDT does reward accurate modeling of these interactions. Additionally, lDDT has recently been updated to exclude rewards for clashing atoms in nucleic acids^45^, thus enabling integration of clashscore and lDDT into one concise score. We hence chose to continue our assessment with lDDT alone, including its penalty for steric clashes.

There was a large variation in performance across targets, but whether a specific target was easy or hard was straightforward to evaluate based on the general range of scores using any of the three chosen metrics (**Figure 2A-C and Supplemental Table 2**). In particular, it was immediately clear that the availability of templates (identified using a sequence-independent method, US-align^34^) was important for target prediction accuracy. While CASP16 groups were able to predict an acceptable fold (TM-score > 0.45^35^) for 21 of 36 targets, only 2 of these did not have a template (template TM-align < 0.45) (**Supplemental Table 3**).

**Figure 2:**
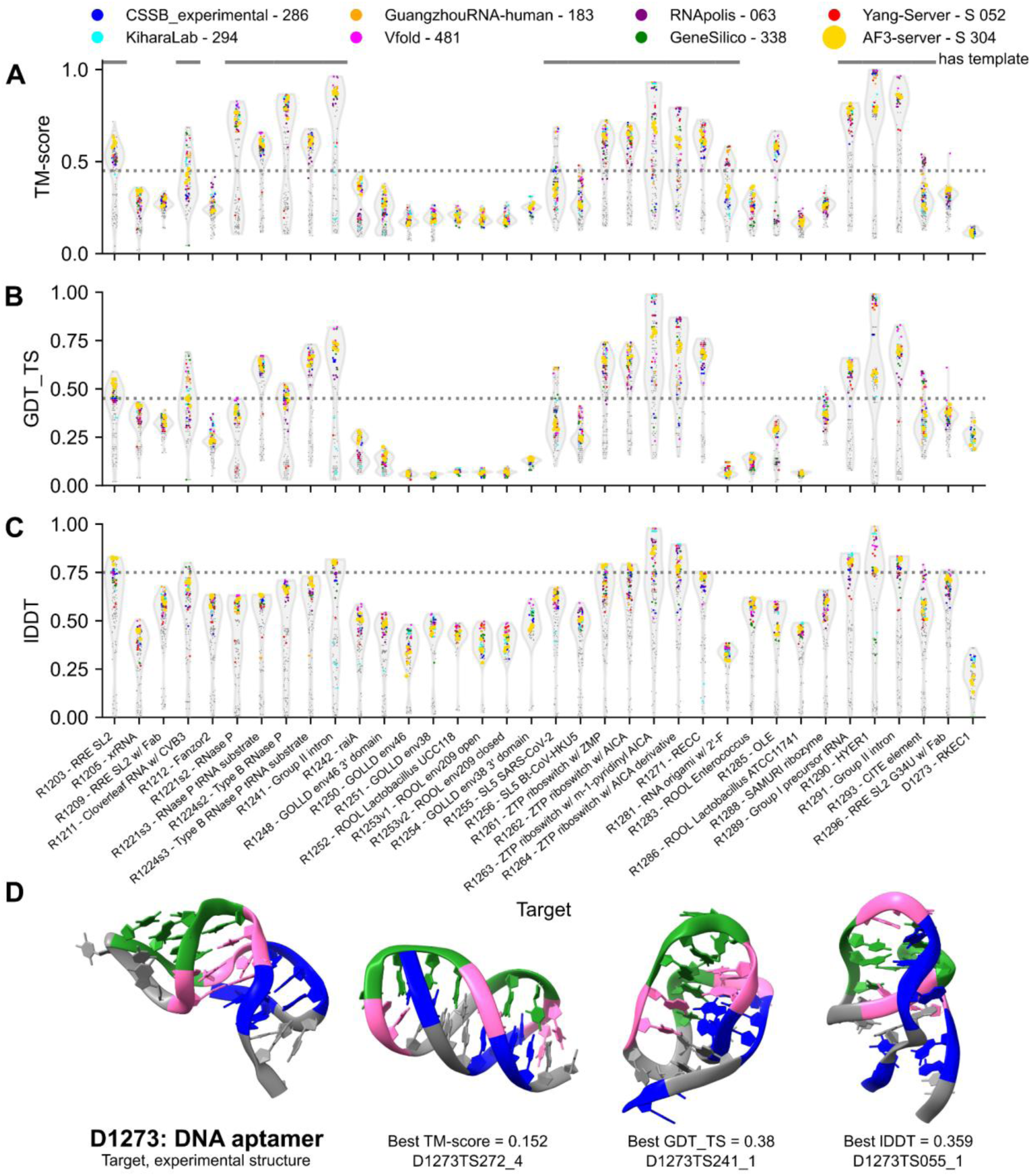
Performance of predictors on nucleic acid monomer assessment targets. For each target, a violin plot is displayed showing the scores of all models for (**A**) TM-score, (**B**) GDT_TS, and (**C**) lDDT. The violin plot’s extent shows the range of scores, while its width represents the density of models, normalized to the maximum density. Models from groups of note are represented by a colored circle, while all other models are gray circles. The threshold for an “acceptable” score, following ^35,71,72^, is labeled with a gray dashed line. (**D**) D1273, a DNA aptamer for dopamine is displayed (left) with the top models by the three scores displayed right. The DNA aptamer is colored by domain: the ligand binding domain in pink and two non-canonical stems on either end in green and blue. The same residues are colored in the predicted models.

Out of all the targets, the DNA aptamer for dopamine, D1273, had the poorest scores by TM-score or lDDT, with no team predicting the correct fold (**Figure 2A,C**). This target’s experimental NMR structure, which included only two Watson-Crick-Franklin base pairs (as supported by the structure ensemble as calculated in AMBER with 872 unambiguous distance restraints, including those based on NOESY cross-peaks involving normally exchangeable nucleobase imino and amino ^1^H nuclei, and including 43 intermolecular NOE cross-peaks; personal communication with structure authors P. Johnson and D. Mackereth), deviates from the secondary structure previously proposed in the literature^68–70^ or in any of the CASP16 predictions (**Figure 2D**), revealing that DNA structures are a significant challenge for existing prediction methods.

### 3.3 | Difficulty and Performance of NA Monomer Predictions

To learn more about performance, we clustered TM-score by group and by target. Across groups, a lack of strong clustering indicated that target difficulty was generally even across the different methodologies used by groups (**Figure 3A**). Many groups sampled structures from several of the same structure prediction methods, which would explain similar performances. The groups that generally had the highest TM-scores were not servers, used a combination of many techniques including deep learning and non-deep learning techniques, and used templates and MSAs (**Figure 3A**). The use of literature to guide modeling was enriched in groups that performed well, suggesting that the integration of knowledge of past experiments – something still mainly conducted by human expert researchers – remains of vital importance.

**Figure 3:**
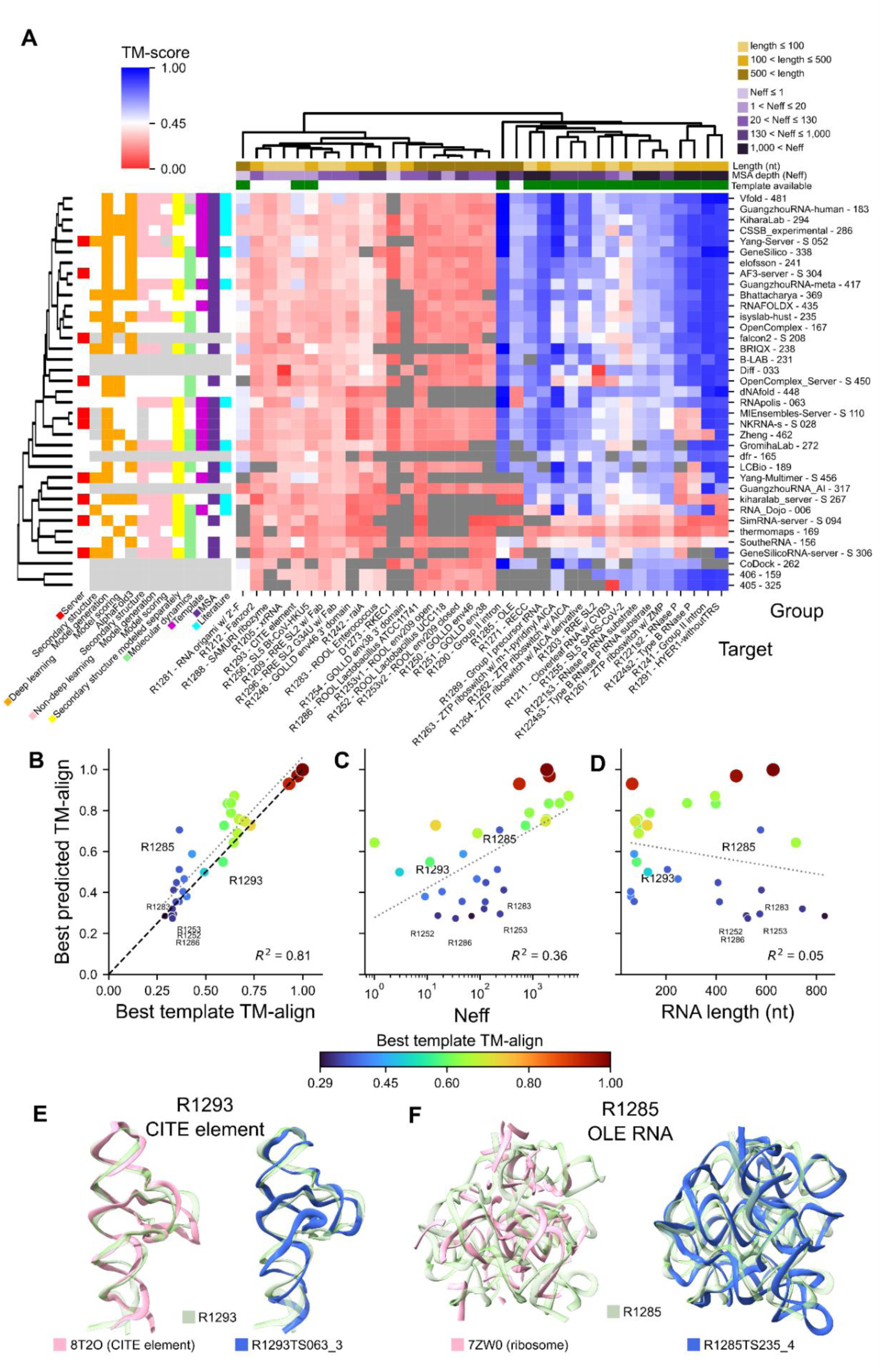
Difficulty of nucleic-acid monomer targets. (**A**) The best predicted TM-score for groups and targets are plotted. Only groups that participated in > 60% of targets are included. The targets are clustered using Euclidean distance of TM-scores, enforcing sequence-dependent alignment. For groups, a tree was created using Euclidean distance as well, but the leaf order optimized to order the groups’ performance. The sum of Z_TM-score_>0 was used to measure performance. The top group was placed at the top, and the tree grew downwards from there. At each node, the branch with the highest average score for its top three teams was added first. If a group did not submit models for a target, the box is colored gray. The groups and targets are categorized on the left and top of the figure, respectively. Groups are labeled as having used a technique if they employed it for any target; for example many groups used AlphaFold 3 just for targets that were too large for other methods. (**B-D**) The best predicted TM-align scores for all RNA targets are plotted against (**B**) TM-align of the best template, (**C**) the normalized number of effective sequences (Neff), and (**D**) RNA length. The targets are colored by TM-align of the best template. In (B), the line where the best template equals the predicted TM-align is labeled in black. For (B-D) the line of best fit is in gray, with the R^2^ value reported at the bottom. Targets of note are labeled. (**E**) R1293, a translation enhancer element, and (**F**) R1285, a OLE RNA, are displayed in transparent green. The best template by TM-align is overlaid in pink on the left and the best prediction by TM-align is overlaid in blue on the right. For OLE, only the nucleotides in 7ZW0 that US-align aligned with the target are displayed.

When clustering across targets, however, we observed two clearly separated clusters (**Figure 3A**). In the first cluster, all groups performed poorly and only 2 of these 18 targets have a template (TM-align > 0.45). Within this template-free cluster, 7 of these 18 targets are large (> 500 nt) and sub-cluster together. In the second cluster, where all groups generally performed well, 17 of 18 of these targets have a template. The one target in this second cluster without a template, R1285 OLE RNA^73^, had a large amount of evolutionary data, a deep MSA, and functional data in the literature^74^ and is further discussed below. Groups that did not use templates in modeling had lower TM-scores for certain targets with templates such as the ZTP riboswitch (R1261, R1262, R1263, R1264) and cloverleaf RNA (R1211) (**Figure 3A**).

Over all targets, we observe a strong dependence of predictor performance on how close an available template was to the target, as parameterized by the sequence-independent TM-align score (**Figure 3B**). MSA depth also explained some variance in predictors’ performance on targets (**Figure 3C**). RNA length was not correlated with predictor performance, although for targets without a template, there appeared to be a trend where predictors received worse TM-align scores for longer targets (**Figure 3D**).

For “easy” targets with a very good template (which we defined as TM-align > 0.8), predictors were able to predict structures with similar accuracy to the template, independent of MSA depth or RNA length. For “medium” targets (template with 0.45 < TM-align < 0.8), we observe some targets where predictors outperformed the template. Notably, however, for R1293, a viral translation enhancer element, the best-predicted TM-align was lower than the template TM-align. This indicates that atomic accuracy in refinement or recognition of a previously available template (PDB 8T2O) was not achieved (**Figure 3E**). The sequence identity of this template to the target was low, 58 %, suggesting the importance of extending future template searches beyond sequence, for example, through the use of covariance models that can be informed by secondary structure models^75^. In the “difficult” target category (best template TM-align < 0.45), better performance was observed for shorter targets and targets with deep MSAs. In particular, there was one outlier, the R1285 OLE RNA noted above, which shows that groups were able to make reasonable predictions for a long target without a template (**Figure 3F**). The closest previously available structure for this target by TM-align score was from the ribosome (PDB 7ZW0, TM-align = 0.365) and is clearly not a viable template (**Figure 3F)**. This target, however, did have a large MSA (Neff = 235), suggesting that predictors may be capable of template-free modeling given sufficient evolutionary data.

### 3.4 | Ranking Group Performance on NA Monomer Targets

As in CASP15^6^, we used a Z-score–based ranking system (see **Methods** and **Supplemental Table 4**) to order groups by performance (**Figure 4A-B**). Vfold unambiguously performed the best, consistently performing well on all targets. Four other teams – GuangzhouRNA-human, KiharaLab, Yang-Server, and GeneSilico – also performed well, with each group performing below average on only a handful of targets. Many teams outperformed the baseline of the AlphaFold 3 server, submitted by the Elofsson group, including one server, Yang-Server. The ranking, in particular the first and top five groups, was robust to alternative choices in scoring methods and score weighting, such as direct use of TM-score or of lDDT instead of Z-scores (**Supplemental Figure 2**).

**Figure 4:**
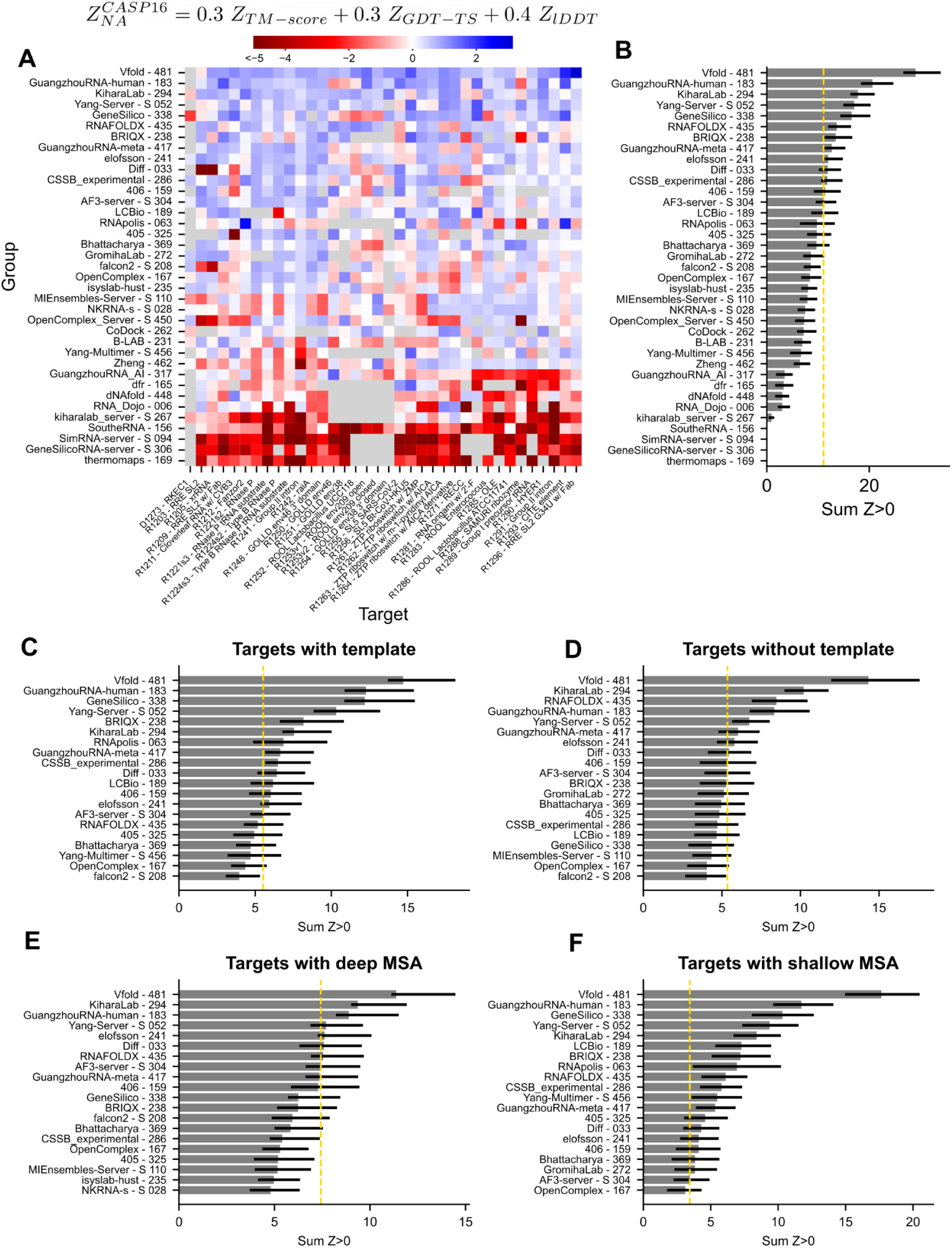
Ranking of groups for the nucleic acid monomer category. (**A**) For every target (columns), the *Z*^*CASP*^^16^ is plotted with blue representing more accurate predictions relative to other groups for that target. Only groups that participated in more than 60% of targets are displayed. (**B**) For every group, their positive *Z*^*CASP*^^16^ scores were summed to obtain a final ranking. The performance of the AlphaFold 3 server, is marked with a gold dotted line. The targets, excluding the DNA target, are then separated based on (**C**) the presence of a template (TM-align > 0.45; N=18) and (**D**) targets without template (N=17); and (**E**) the presence of a deep MSA (Neff > 130; N = 19) and (**F**) targets with a shallow MSA (N=16). Only the top 20 groups for each category are displayed.

The targets can be separated into template-based modeling challenges (targets with a known structure with TM-align > 0.45) and template-free modeling (**Figure 4C-D**). Vfold ranks top in both categories. Amongst the other top five teams, GuangzhouRNA-human and GeneSilico rank better for template-based modeling, and KiharaLab ranks better for template-free modeling. We noted that for template-based modeling, the AlphaFold 3 server ranked poorly, indicating that human predictors, and even the automated Yang-Server, may be better able to use template information; indeed, AlphaFold 3 does not carry out template search for nucleic acids^20^. For the template-free modeling, an additional group, RNAFOLDX, performed well, though still behind Vfold and KiharaLab.

Additionally, Vfold performed uniquely well on targets without a deep MSA (Neff < 130) (**Figure 4E-F**). We note that some of these targets did have functional data, experimental secondary structure, or larger MSAs in the literature that were not identified in the automated rMSA. Hence, automated approaches may focus on improving methodology for obtaining or better interpreting MSAs to bridge the gap between human and automated approaches^11^. Supporting this picture, the AlphaFold 3 server ranked better (8th vs. 19th) on targets with deep MSAs compared to shallow MSAs.

### 3.5 | Base Pair Prediction Accuracy

Complex and novel targets appear well beyond current capabilities for NA 3D structure prediction. However, RNA folding can be simplified into a hierarchical process^76^: secondary structure – the pattern of canonical base pairs – forms a set of RNA stems which are then stitched into the overall 3D fold. After CASP15, retrospective analyses suggested that the difficulty in predicting secondary structure contributed to poor accuracy in RNA modeling, particularly by automated methods^30,77^. CASP16 offered the prospect of carrying out tests of secondary structure accuracy prospectively. The secondary structure of all targets, here defined as the list of all Watson-Crick-Franklin and Wobble pairs, turned out to be predicted to a high level of accuracy (**Supplemental Figure 3A**). In fact, CASP16 predictors, including automated methods, outperformed widely used algorithms that predict secondary structure without explicit 3D modeling or MSA input (**Supplemental Figure 3A**). Furthermore, unlike 3D structure prediction, secondary structure prediction accuracy was only weakly correlated to factors that can mitigate 3D structure prediction difficulty such as the availability of template and a larger MSA (green lines in **Figure 5A**). The trend in RNA secondary structure performance is more reminiscent of the performance observed in current protein 3D structure prediction^78^, suggesting these prediction algorithms are reaching sufficient accuracy in their prediction of secondary structure to be important and useful in structural research.

**Figure 5:**
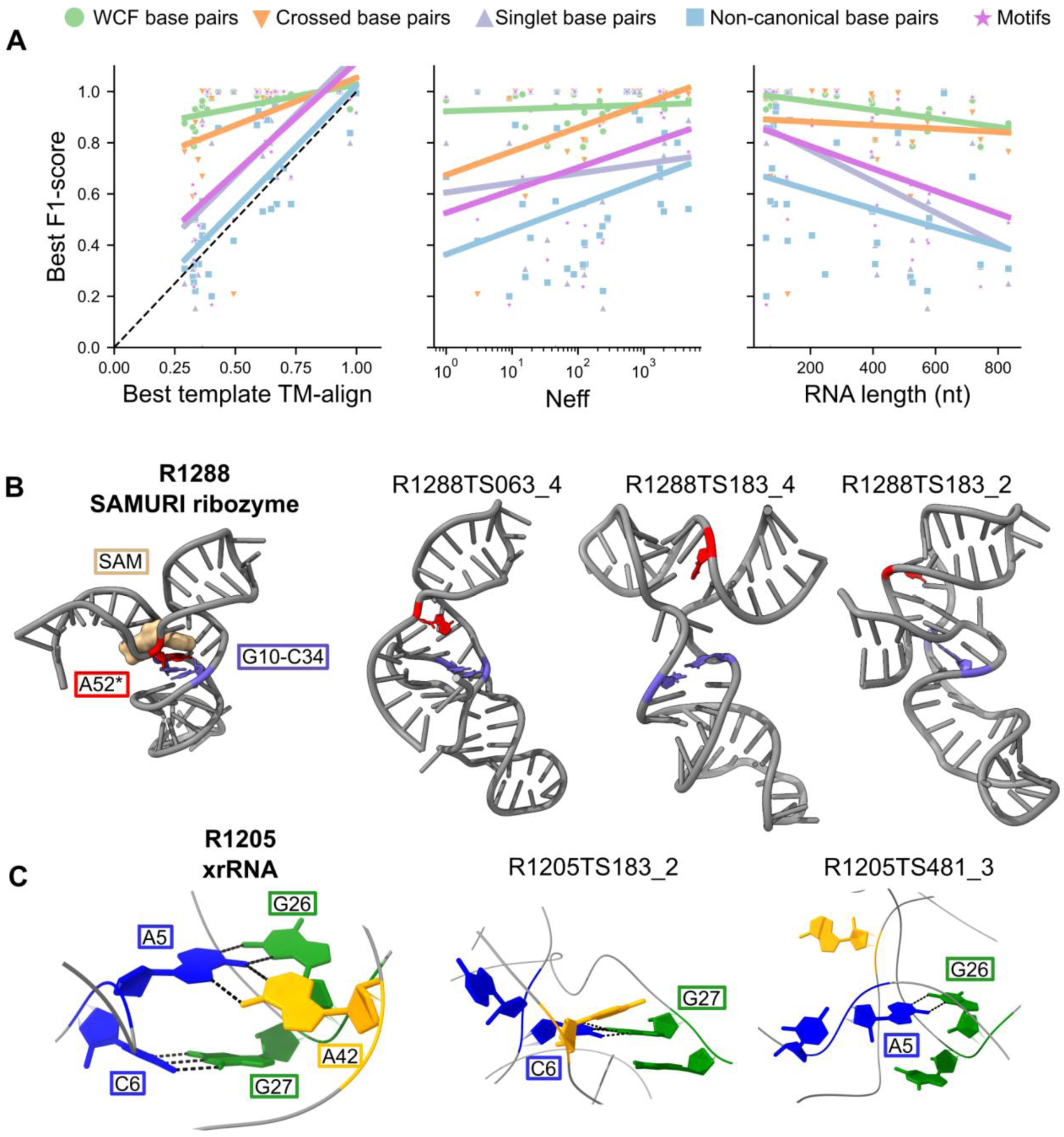
Base pair and motif prediction accuracy. (**A**) For each target, the best F1-score across all groups is plotted for (green) all base pairs, (orange) crossed base pairs, (purple) singlet base pairs, (blue) non-canonical base pairs, and (pink) motifs. Targets that do not contain the interaction are not plotted. These best F1-scores are plotted against the best template TM-align, MSA Neff, and RNA length. A linear fit is plotted for each interaction type. (**B**) R1288 (SAMURI ribozyme) is displayed with the singlet base pair highlighted in purple, the reactive adenine in red, and the ligand in brown. Three predictions that contain the singlet base pair are displayed; none of these predictions correctly modeled the catalytic active site. (**C**) The pseudoknot crossed pair and triplet interactions for R1205 (plant virus xrRNA) are displayed. The models that predicted one pair in this interaction are displayed. Hydrogen bonds are displayed as black dotted lines.

Despite these positive indicators of secondary structure prediction accuracy, performance was more modest for subsets of RNA base pairs that are hallmarks of complex tertiary structures. Some targets contained pseudoknots, which are sets of base pairs in which residues in a stem’s loop base pair with residues outside the loop. These pseudoknots encode pairs that ‘cross’ each other and hence are not nested like conventional RNA secondary structures. Some targets also had singlets, which we defined as base pairs not involved in a longer stem of canonical pairs but are instead usually stabilized by interactions with other residues in non-canonical structural motifs. These crossed and singlet base pairs can play an integral role in defining the 3D fold of an RNA by bringing distal regions of sequence together, but secondary structure algorithms often do not predict or have poor accuracy in predicting these pairs. The CASP16 results showed that accuracy in predicting crossed and singlet base pairs was poorer than base pairs generally (**Supplemental Figure 3B-C**). Singlet prediction performance depended on the availability of a template and RNA length (purple lines in **Figure 5A**). For crossed base pairs, the performance of predictors was not dependent on the length of the RNA but was dependent on MSA depth (orange lines in **Figure 5A**). This result suggested that the predictors were able to extract the signal for crossed pairs from MSAs, whereas the singlet information was more difficult to extract from, or was not contained in, MSAs.

Vfold, GuangzhouRNA-human, RNApolis, and GeneSilico stood out in their ability to predict crossed base pairs accurately for difficult targets like the translation enhancer element, R1293 (**Supplemental Figure 3B**). Another example, R1288, is an *in vitro* selected ribozyme and hence does not have a template or MSA. This SAMURI ribozyme binds S-adenosylmethionine (SAM) and catalyzes the alkylation of nucleotides^79,80^; A52 is the alkylated nucleotide in this target. Around the binding site, immediately upstream of the second stem, G10 crosses around the first two base pairs of this stem to form a singlet base pair with C34. This crossed singlet base pair is important in forming the ligand pocket, sandwiching the adenine that will be modified with the SAM cofactor. While most of the secondary structure was previously proposed in the literature^79^, the structure of the binding site and this singlet base pair was not known during CASP16 prediction. Only GuangzhouRNA-human and RNApolis predicted this singlet base pair, in addition to an algorithm that predicted secondary structure without explicit 3D modeling, RibonanzaNet (**Supplemental Figure 3B-C**). Despite predicting this important singlet base pair, the groups did not accurately predict the 3D structure of the active site (**Figure 5B**). GuangzhouRNA-human predicted A52 stacked between bases at the junction, far from the target binding site. Vfold did correctly predict A52 bulged out but predicted an extended binding site that would be unlikely to be able to restrain the reactive nucleotide sufficiently for catalysis.

R1205 is a class 3 exoribonuclease-resistant RNA (xrRNA) which is known to form a conformational ensemble of open and closed structures; the closed structure is the active conformation and contains an additional pseudoknot^81^. There was no previous structure of the closed, active conformation. Predictors were not told it was the active conformation, but they were able to submit up to five models and hence could have submitted structures in open and closed conformations. One of the crossed singlet pairs (C6-G27) forms the base of the core of the xrRNA (**Figure 5C**). Of all the predictors, GuangzhouRNA-human and GuangzhouRNA-meta were the only groups to reach a high level of accuracy in predicted crossed and singlet pairs, including the C6-G27 pair (**Supplemental Figure 3B-C**). However, this base pair is not conserved and only weakly affects the activity of the RNA^81^. Instead, the non-canonical triplet base interactions of A5-G26 and A5-A42 appear to be the conserved and functionally important interactions^81^, suggesting the need to analyze pairs beyond canonical base pairs. Vfold predicted the A5-G26 base pair but no group predicted the full triplet interaction of A5-G26 and A5-A42 (**Figure 5C**). Across all targets, Vfold and GuangzhouRNA-human predicted non-canonical base pairs the best. However, with an average F1-score less than 0.5, non-canonical base pair prediction remains a challenge (**Supplemental Figure 3D**).

Base pair prediction accuracy did not clearly discriminate between prediction groups, with many groups performing roughly equivalent overall (**Supplemental Figure 4**). However, the top four in the overall ranking, GuangzhouRNA-human, Vfold, GuangzhouRNA-meta, and GeneSilico stood out from other groups for targets with shallow MSAs (**Supplemental Figure 4E**).

### 3.6 | RNA Motif Prediction Accuracy

Crossed, singlet, and non-canonical base pairs are a subset of the interactions that stitch helices together. RNA motifs beyond base pairing include A-minor interactions, U-turns, T-loops, and more. These interactions can define the 3D fold of a molecule by gluing distal regions together or creating distinct kinks and turns in helices.

Like base pairing, many groups performed roughly equivalently in motif prediction. Interestingly, however, for motifs, automated methods were able to match the performance of human groups, with the AlphaFold 3 server and Yang-Server performing within the error of the top-ranking group (**Supplemental Figure 5**).

Some motifs like tetraloop receptors, Z-turns, and Loop E submotifs were predicted with reasonable accuracy. Motifs like Loop E submotifs have a clear sequence conservation signature, so it was unsurprising that human experts and automated methods could identify these. However, prediction accuracy was poor for other tertiary motifs, potentially because these interactions do not often have obvious sequence motifs or established evolutionary signals. Some particularly challenging motifs were UA-handles, platforms, and A-minor interactions. Predictors were more accurate at predicting tetraloop receptors and T-loops than the submotifs contained within them, a dinucleotide platform and a UA-handle, respectively, which can also arise in other contexts. This result suggests that these larger motifs are easier to predict, potentially because their sequence or evolutionary signal is more recognizable. For T-loops, human predictors, especially KiharaLab and GeneSilico, significantly outperformed automated methods, suggesting there is expert knowledge about the identification or refinement of T-loops that automated methods have yet to learn.

Across RNA structures, there are two particularly common, but challenging, tertiary interactions to predict: the intercalated T-loop and the A-minor interaction. In these interactions, a nucleotide, often distal in sequence and secondary structure, intercalates into a T-loop or docks into the minor groove of a helix, respectively. Groups were able to predict half of the interactions, for example the T-loop motif, with good accuracy. However, predictors were not able to predict which nucleotides intercalate or dock into these positions. For A-minor interactions, there were cases where groups correctly predicted the identity of adenines that were docking into minor grooves, but did not dock them into the correct helix (as reflected in the higher F1 scores for docked-A than A-minor).

Returning to xrRNA target R1205, the previously described A42-A5-G26 triplet is connected to a larger motif, a T-loop (residues 24-29 and 31) with an intercalated A (A5). No group predicted the T-loop would fold, despite homology to xrRNA molecules from other plant viruses with known structures (e.g., PDB 6D3P), suggesting that template searching, even by human experts, requires improvement. In the previously available templates, the intercalating A was from a different chain in the crystal structure than the T-loop. Hence, inter-chain contacts may need to be more carefully handled in template searches or in preparing training datasets for machine learning.

These results indicate that automated approaches can recognize some of the patterns that govern RNA structural motifs. However, some interactions, like the A-minor interactions, remain challenging, and the stronger performance of the human groups indicates there is room for automated methods to improve. In these analyses, there were a handful of interactions that were predicted by CASP16 groups but were not in the experimentally solved monomer structure but are actually part of the quaternary structure; that is they involved base pairing between two chains of RNA. Differentiation between tertiary and quaternary contacts is a new and challenging task for predictors that we assessed in the RNA multimer category, discussed next.

### 3.7 | Assessment of NA-NA Interfaces and RNA-Only Multimer Targets

As a new category in CASP16, and a new challenge for RNA structure prediction generally, there were 11 RNA-only targets that formed homo-multimers. These targets were released to predictors in two rounds. In round 0, predictors were told that the RNA formed a homo-multimer, but they had to predict the stoichiometry. In round 1, the stoichiometry was provided. The results from round 0 demonstrated that predictors were not able to accurately predict the oligomerization state of the targets (**Supplemental Figure 6A**). Even after stoichiometry was known in round 1, predictors were not able to predict the symmetry correctly, with predictions dominantly predicting no symmetry or the highest order cyclic symmetry even for the many targets that instead had dihedral symmetry (**Supplemental Figure 6B**).

For round 1, 32 of the 65 groups submitted multimeric predictions resulting in 1,072 total predictions, ∼10% of which predicted no meaningful interactions between chains, that is the chains were totally overlapping or the chains were totally separated, with no contacts formed at all. RNA-RNA interchain interfaces are generally composed of the same interactions as tertiary motifs, for example base pairing between two chains of RNA, so we conducted a similar analysis of base pairs and motifs as carried out above for RNA monomers. Groups correctly predicted the kissing loop in R1290 (group II intron), for which there were template structures with perfect sequence identity (**Supplemental Figure 7A**). Additionally, predictors were able to predict some of the interface interactions in R1285 (OLE RNA) (**Supplemental Figure 7A**), but, as discussed elsewhere in this issue, a maximum of 1 interface was predicted in any model^21^. Many of the RNA homo-multimer targets contained intermolecular A-minor interactions. However, groups were only able to predict the correct adenines underlying these interactions in a few cases and they were unable to predict where the adenines docked (**Supplemental Figure 7B**). For example, in R1251 (GOLLD env-38 RNA), A34 and A35 docked in the minor groove of a helix in another chain forming two intermolecular A-minor interactions (**Figure 6A**). Two models (from GuangzhouRNA-human and –meta) predicted both A34 and A35 would dock, and seven other models predicted only one would dock (from CoDock, GuangzhouRNA-meta, and the AlphaFold 3 server). None of the predictions docked the adenines into the correct helix, and only three of these models (the AlphaFold 3 server, GuangzhouRNA-human and –meta) predicted these adenines would dock into a helix from another chain. Qualitatively, these models also had many inter-chain clashes, a common occurrence in the RNA multimer category, creating unrealistic interfaces and low precision of predictions.

**Figure 6:**
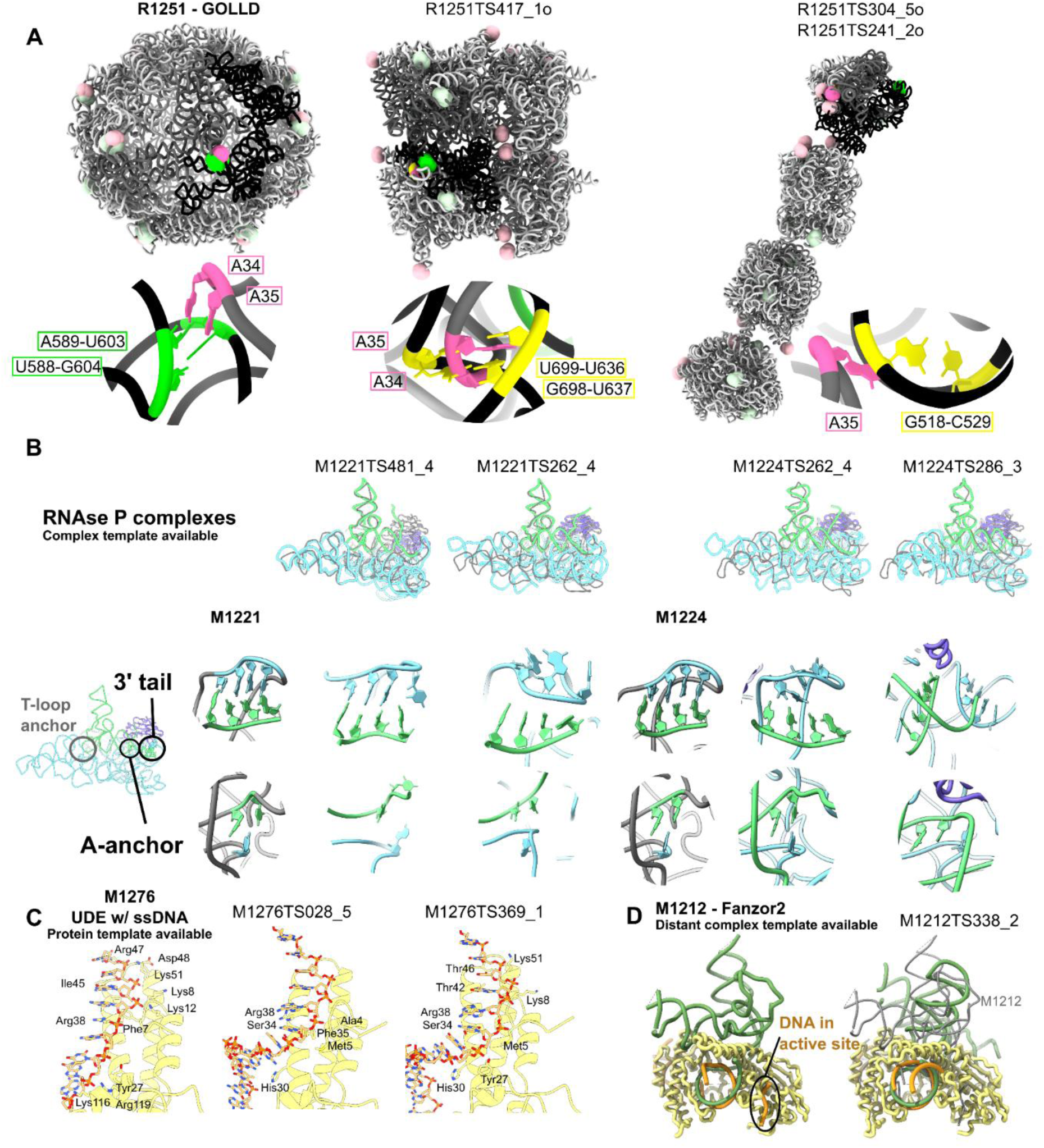
Highlights and problems in nucleic acid interface prediction. (**A**) R1251 (GOLLD env-46 RNA) reference structure and two predicted models are displayed. The target and predictions are composed of 14 chains, a pair of interacting chains colored dark gray and black with the other 12 chains in light gray. In the reference intermolecular A-minor interactions, the adenine (pink) and its base pair docking site (green) are labeled in bright colors for the two interacting chains and lightly colored in other chains. For each prediction, the base pair the adenine is docked into is labeled in yellow. (**B**) The best NA-NA i-lDDT and the best NA-NA ICS models are displayed for the RNAse P complexes M1221 and M1224 with the experimental reference structure in gray, protein in purple, tRNA in green and RNAse P ribozyme in blue. Below, the regions of the 3’ tail base pairs and A-anchor are displayed under each prediction with the experimental structure on the left. (**C**) The interface between protein (yellow) and DNA (orange) in M1276 (cryptic DNA-binding protein UDE) is displayed. The protein residues within 3.2 Å of the DNA are displayed and labeled. (**D**) The protein (yellow), DNA (orange) and RNA (green) of M1212 (Fanzor2 complex) are displayed with the target on the left and the best prediction by NA-NA and NA-protein ICS on the right superimposed on the gray target DNA and RNA.

To assess complex prediction overall, as with monomer metrics, we used metrics that could be applied across all biomolecules, which we split into two categories. First, we assessed the global 3D fold and local accuracy of the entire complex using the same metrics as the NA monomer category, GDT_TS, TM-score, and lDDT (**Supplemental Figure 7C**). Second, we scored the interface between monomers using scores previously used in CASP protein multimer categories, ICS, IPS, and i-lDDT^42^ (**Supplemental Figure 7D**). Predictions of R1290, a dimeric RNA target with an excellent template, confirmed that excellent-quality RNA-RNA interfaces received high scores for these metrics. However, interfaces for no other RNA multimer were predicted accurately. Even a simple interface as in R1281, a kissing loop connecting two six-helix bundle origamis captured in a dimer conformation, was poorly predicted. While we can rank the prediction methods, prediction quality across all methods was too poor to be considered reliable (**Supplemental Figure 7E**).

Further analysis of intermolecular interactions revealed an intriguing trend. There are four targets representative of the ROOL RNA family: R1252, R1253, R1283, and R1286. The ROOL family had no known structure prior to CASP16, and the targets were not homologous to any RNA in the PDB. Despite originating from the same RNA family with the same literature sequence alignments and biochemical data, predictors submitted more accurate predictions for R1283, the target with the deepest MSA. The top TM-score for R1283 was 0.376, compared to 0.284 for R1252, which had the shallowest MSA (**Figure 3C**). In addition, the interfaces for R1283 predictions were substantially more accurate, with an ICS of 0.286 compared to an ICS of 0.071 for R1252. These targets all had similar folds, lengths, and templates, enabling a more systematic exploration of other variables that impacted prediction accuracy. A standard MSA pipeline (rMSA) obtained a deeper MSA (551 sequences, Neff = 281) for R1283 than the other ROOL sequences; the shallowest MSA resulted from R1252 (43 sequences and Neff = 16). Prior data showed that these ROOL sequences can be integrated into the same large MSA^82^, suggesting that the standard automated MSA generation program may benefit from improvement. Due to the large size of these complexes, many groups resorted to using AlphaFold 3 to obtain initial models, and so the performance of AlphaFold 3 on these targets could impact overall performance. The AlphaFold 3 server predicted a more accurate structure for R1283 than the other ROOL sequences; this result was replicated post-CASP season for AlphaFold 3, both with and without MSA. Whether the better performance of AlphaFold 3 was due to a deeper MSA or other factors should be investigated further to systematically understand where the prediction faults lie.

Additionally, predictions were more accurate for the tetrameric state of this complex (R1283v2, TM-score = 0.376, ICS = 0.286) than the octameric state (R1283v3, TM-score = 0.328, ICS = 0.206). The interface between the tetramer has more interactions and a stronger evolutionary signal than the interface that glues two tetramers together^73^, which may explain these trends and support the ability of algorithms to extract structural data with sufficiently deep MSAs.

Beyond the RNA-only homomeric multimers described above, RNA-RNA interactions were also observed in hybrid targets, defined here as the targets containing both NA and protein. Predictors were able to predict simple double helix interfaces with perfect complementarity between strands (**Supplemental Figure 7F**). However, accuracy was much lower for interactions more complex than double helices, even when good templates existed. M1221 and M1224 are each a complex of an RNAse P ribozyme, a precursor tRNA, and the RNAse P protein. RNAse P must recognize the correct tRNA and position the 5’ leader correctly for scission accomplished through 3 major interaction sites: the T-loop anchor, the A-anchor, and a set of base pairs involving the 3’ tRNA tail^83^ (**Figure 6B**). These RNA-RNA interactions are well-established in the literature, so despite no sequence-identical templates, the modest accuracy of RNA-RNA interface scores was surprising. The top models by i-lDDT and ICS, provided by Vfold, CoDock, and CSSB_experimental, all had accurate global folds. In particular, the RNAse P-tRNA interfaces aligned well with the reference cryo-EM structures. However, the models lacked accuracy at the A-anchor and 3’ tail sites. RNAse P has a complementary sequence to the 3’ tRNA tail ACCA to tether it in place, which Vfold modeled well. CoDock and CSSB_experimental did not model these base pairs. The A-anchor is an adenine that extends from a bend in RNAse P to pair with the base in tRNA immediately upstream of the scission site, positioning the tRNA backbone for the reaction. All four of these models had the adenine correctly in a bulged sequence, and CoDock and CSSB_experimental had the bases surrounding the scission site in the tRNA stacked as in the targets. Despite accuracy in the individual structures, only CoDock in target M1224 positioned the chains accurately relative to each other. It appears that RNA-RNA interface prediction is challenging even for well-studied systems; predictors were able to model the individual components from templates, but had difficulty in assembling them in a complex.

### 3.8 | Assessment of NA-Protein Hybrid Targets

Using our biomolecule-agnostic metrics, the hybrid targets could be scored similarly to the RNA multimer complexes. To emphasize the NA predictions, we focused on the NA-NA and NA-protein interfaces, excluding the protein-protein interfaces, which are evaluated in detail elsewhere in this issue^22^.

In round 0, the stoichiometries of two DNA-protein targets were predicted with 60% and 31% accuracy after removing predictions that did not model any nucleic acid (**Supplemental Figure 6C**). This suggests that even though DNA-protein complexes had much prior structural work, unlike RNA multimers, stoichiometry prediction is still hard. For example, in M0287, the majority of predictors modeled only one DNA chain when the DNA should be in duplex form, making assessment of the interactions hard to interpret. Hence, we focused on round 1 where stoichiometry was provided to predictors.

Highly accurate interfaces were only predicted for three complexes: M1209, M1293, and M1296 (**Supplemental Figure 8A-B**), which all were Fab-RNA loop interactions commonly used to crystallize RNA^84^. A simple target, M1276, a short single-stranded DNA binding protein, demonstrated that these scores are robust for DNA as well. M1276 predictions had high ICS and IPS scores but lower i-lDDT scores, indicating that predictors found the correct interacting residues but did not model the interaction correctly. Indeed, when we examined the protein residues within 3.2 Å of the DNA strand, there were key missing features in the predictions, including missing polar residues along the backbone (**Figure 6C**). In addition, all models missed an intercalating arginine, which perturbed the local DNA conformation.

While performance appeared better for NA-protein interfaces than for RNA-RNA interfaces, all the RNA multimers except one had no 3D structure templates. Similar to poor performance on RNA-RNA interfaces,, predictors were unable to identify residues involved in the interfaces of NA-protein targets without templates, M1282 and M1211. Scores for RNA-RNA interfaces for the previously described M1221 and M1224 were similar to the RNA-protein interface scores for these targets (**Supplemental Figure 7F and Supplemental Figure 8A**), suggesting predictions of NA-NA and NA-protein interfaces remain inaccurate if no template is available.

The generality of the assessment metrics also enabled the analysis of RNA-DNA-protein complexes. For example, Fanzor2, an RNA-guided DNA endonuclease, is homologous to a previously solved Fanzor1 complex in the NA-protein interface^85^. Predictors generally were able to recognize the homology and model a pseudoknot analogous to the one found in Fanzor1 (**Supplemental Figure 3B**). However, the homology modeling was not refined to atomic accuracy. All predictors missed the NA-protein interaction in the catalytic site of Fanzor2 (**Figure 6D**). Many predictions showed a stem-loop interacting with this protein domain, homologous to Fanzor1, but experimentally the Fanzor2 RNA was not found to have this interaction. Instead, 4 DNA residues were resolved at the Fanzor2 protein active site; the enzyme is likely in an inhibited state^86^, so although predictors did not predict this interaction, it raises the question as to whether this conformation is relevant functionally and could have been captured within the five submissions made by each CASP16 group.

To emphasize NA modeling, our rankings place emphasis on NA-containing interface accuracy. With this approach, KiharaLab ranks the best, followed by CSSB_experimental, Vfold, and Zheng, with MIEnsembles-Server as the top-ranking server (**Supplemental Figure 8C**).

### 3.9 | NA-Ligand Targets

RNA and DNA aptamers that bind small molecule ligands are functionally important molecules for which there is rich experimental structure knowledge. However, quantitative assessment of blind predictions of NA-ligand interactions has not been carried out before. Here, the availability of five NA-ligand CASP16 targets and the development of lDDT as a universal metric across proteins, nucleic acids, and small molecules enabled such an assessment. We used the lDDT of the ligand pocket and the lDDT of the NA-ligand interface (i-lDDT) to assess the accuracy of ligand poses (**Supplemental Figure 9**). Groups were able to predict ligands well for a set of three ZTP-riboswitches (R1261, R1263, and R1264) with various ligands, with particularly accurate predictions from CoDock and GeneSilico. All had excellent templates for the RNA structures and ligand poses. However, for the other two nucleic acid ligand targets, a DNA aptamer (D1273) and an RNA enzyme (R1288), no group predicted the nucleic acid structure well and therefore could not accurately model the ligand pocket. Although the state of the NA-ligand predictions field could not be assessed comprehensively due to the limited number of targets, this analysis holds promise for future assessments with a wider range of NA-ligand targets.

### 3.10 | NA Structure Prediction Current State and Historical Perspective

We combined the scores from all four categories assessed herein to obtain a general ranking of available methods for nucleic acid structure prediction (**Figure 7A**). Vfold ranked the highest, followed by Kihara lab. In third place, GuangzhouRNA-human exhibited excellent performance, particularly in the RNA targets, and GeneSilico and CSSB_experimental performed reasonably well in all categories. Despite not participating in ligand or hybrid categories, Yang-server was the next top performer, followed by other groups who participated in more nucleic acid categories. The AlphaFold 3 server was outperformed by 8 groups in the overall ranking and by 12 groups in the NA-monomer category.

**Figure 7:**
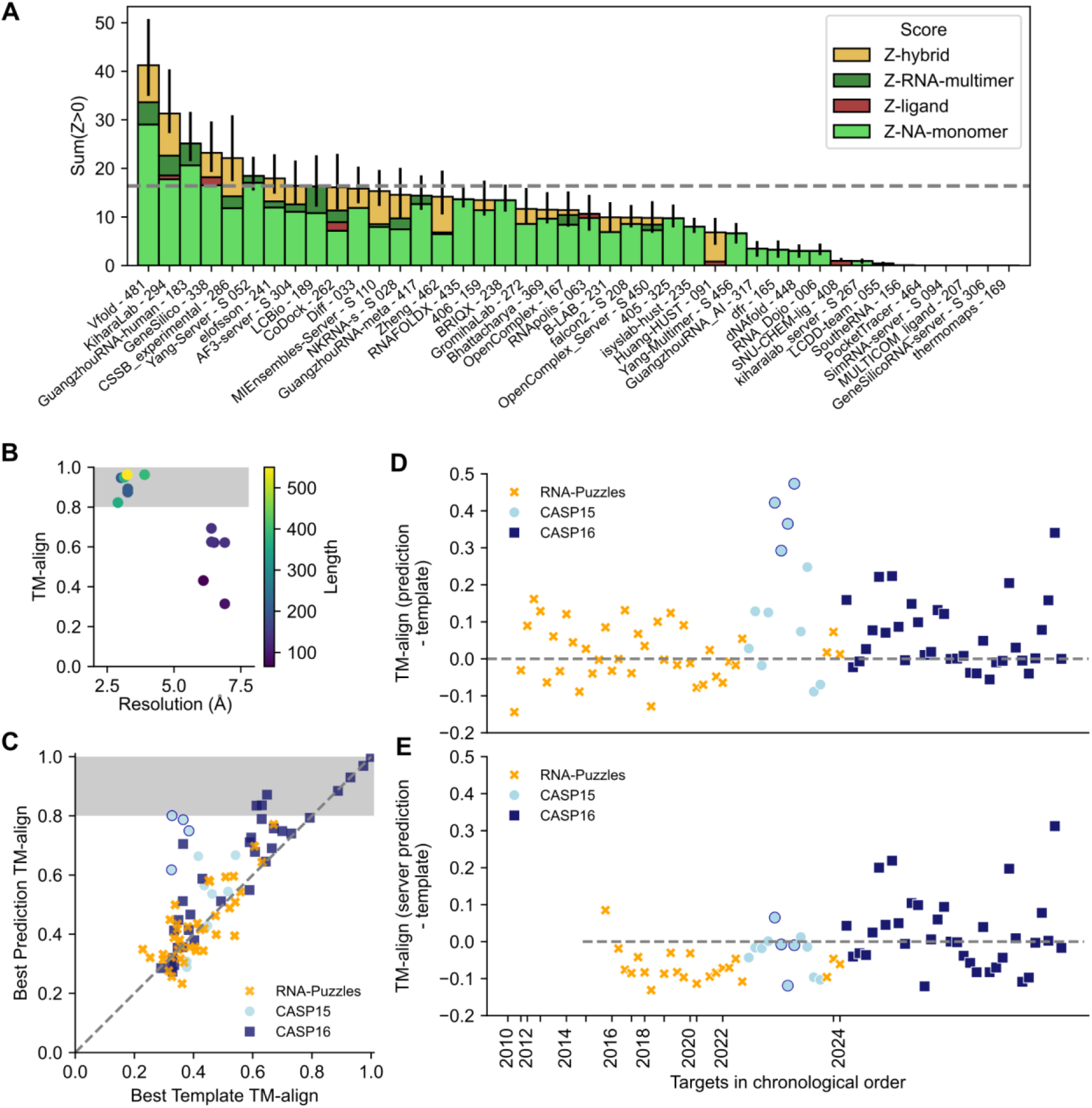
Summary of performance in nucleic acid structure prediction in CASP16 and through time. (**A**) CASP16 ranking for nucleic acid targets. All Z-scores from the four categories are summed. The 68.2% confidence intervals are also summed and displayed. The AlphaFold 3 server performance is displayed with a gray dotted line. (**B**) To estimate experimental error, models of the same RNA sequence solved under distinct experimental conditions by different research groups were compared. The TM-align score between these independently determined structures is plotted against the lowest resolution of the experimental structures. At 2–5 Å resolution, TM-align scores lie between 0.8 and 1.0 (shaded gray box), suggesting that this is a reasonable range of maximal TM-align values given experimental precision. (**C**) The best predicted TM-align is plotted against the TM-align of the best template for all assessed targets from RNA-puzzles (orange X), CASP15 (light blue circle), CASP16 (dark blue square). Human designed, RNA origami targets are circled in dark blue. (**D**) The difference between the best predicted TM-align and the best template TM-align displays how predictors performed relative to the template. All targets are plotted in chronological order. (**E**) As in (D) but only server predictions are considered.

The performance of methods for NA molecules and their complexes appeared generally worse than for the protein molecules and protein-protein complexes, as judged by metrics that could be computed across both types of biomolecules like TM-score, GDT_TS, and lDDT. In principle, the worse scores for NA targets might be explained by greater inherent flexibility and/or poorer experimental precision for RNA and DNA molecules. To test this explanation, we estimated a skyline for NA structure prediction by identifying pairs of RNA structures that have been solved by independent groups at around the same time in recent years – including ROOL, raiA, and OLE RNA molecules that were targets in CASP16 (**Supplemental Table 5**) – and calculating pairwise structural similarity as TM-align scores. For pairs of structures solved with resolution worse than 5 Å, we observed TM-align scores at or above 0.45, the conventional cutoff for two RNAs sharing global folds^35^ (**Figure 7B**). For structures solved at a resolution of 2–5 Å, we observed TM-align scores between independently determined structures in the range of 0.8 to 1.0 (**Figure 7B**). We therefore propose that a threshold of 0.8 should be a reasonable goal for the prediction of biologically well-defined RNA structures, most of which are now experimentally solvable at 2–5 Å resolution. After excluding targets with a previously available template with TM-align > 0.8, CASP16 blind predictions for only three targets surpassed this threshold: the two RNAse P ribozymes and a group I ribozyme precursor tRNA structure (R1221s2, R1224s2, and R1289) (**Figure 7C**). These all still had reasonable templates (TM-align > 0.6), deep MSAs (Neff > 2,000), and extensive structure-function information in the literature; indeed these represent the first two classes of RNA enzymes discovered in nature. This analysis suggests that predictors were able to refine a reasonable template using evolutionary and/or functional information, a promising trend for the future of template-based modeling. However, the prediction accuracy for the other RNA structures, including ones where independently solved structures gave TM-align values above 0.95 (**Supplemental Table 5**), has not approached the threshold of TM-align > 0.8 (**Figure 7C**). Taken together, these results indicate a substantial performance gap between current RNA structure prediction accuracy and the limit set by experimental precision for most targets.

While it was disappointing that the accuracy in NA structure prediction has not improved to the level of protein structure prediction, we wished to understand whether there has nevertheless been measurable progress in CASP16 compared to previous blind challenges. Such an analysis requires calibrating the difficulty of blind prediction targets across history. We noted here that prediction performance was highly correlated to the quality of the best template available, suggesting that the best TM-align across all previously available structures would be a reasonable measure of target difficulty for RNA (**Figure 7C**), analogous to CASP’s historical comparisons for proteins calibrated by best templates^87^. We used this observation to compare across the history of RNA structure prediction blind challenges by tabulating and plotting the difference between the best predicted TM-align and the best template TM-align across RNA-puzzles, CASP15, and CASP16 (**Figure 7D**). There were many targets that were predicted with equal or worse TM-align than the template, in addition to some targets that had up to a roughly 50% improvement from the template. Among the latter, there were five outliers. Four outliers from CASP15 were human-designed RNA nanostructures which did not have templates but for which human expert predictors were able to guess the design objectives from the sequence (circled in dark blue in **Figure 7D**). The other outlier is R1285 from CASP16, OLE RNA, which had no reasonable template but a deep MSA. As MSAs become more important for RNA structure prediction, MSA depth should be included in the assessment of target difficulty.

Even while including the R1285 outlier, we did not observe an appreciable improvement in prediction quality in CASP16 compared to previous years (**Figure 7D**). There were and still remain blind targets where the best TM-align score to a previously available 3D structure template is better than the score for the best blind prediction. If we exclude the four human-designed RNA targets in CASP15, which turned out to be easy for some human predictors, the distribution of improvement of TM-align scores over best available templates has remained similar, with a small but statistically insignificant improvement in the mean value from 0.027 ± 0.014 to 0.060 ± 0.016 (*p =* 0.13 by t-test; **Table 2**). One notable improvement, however, has occurred. Historically, server groups fared poorly in prior blind challenges, nearly always giving worse predictions than human experts and even available structural templates^1–6^. In CASP16, some server predictions, notably those from Yang-Server, have at least improved upon templates (**Figure 7E**). The mean improvement of TM-align scores over best available templates for servers increased from –0.067 ± 0.012 before CASP16 to 0.022 ± 0.017 in CASP16 (*p* = 7.7 x 10^−5^; **Table 2**). While still inferior to human-guided modeling, automated modeling approaches will likely play an increasingly important role in the NA structure prediction community. Additionally, human groups appear to be increasingly taking advantage of automated approaches, especially the AlphaFold 3 server, to sample more diverse ensembles of models and then manually curate the best predictions, as evidenced by the methodologies reported by CASP16 top-ranking teams.

**Table 2:** Comparison of prediction accuracy prior to CASP16 and in CASP16.

| Predictors | Number of targets considered<sup>†</sup> | Time of prediction | Best predicted TM-align improvement relative to best template TM-align | | $p$ (t test) |
| --- | --- | --- | --- | --- | --- |
|  |  |  | Mean | SEM |  |
| All | 31 | Pre-CASP16 | 0.027 | 0.014 | 0.13 |
| All | 34 | CASP16 | 0.060 | 0.016 |  |
| Server | 31 | Pre-CASP16 | -0.067 | 0.012 | 0.000077 |
| Server | 34 | CASP16 | 0.022 | 0.017 |  |

<sup>†</sup> Human-designed targets and targets for which no server participated have been excluded. SEM: standard error of the mean.

## Discussion

CASP16 provided a wide range of prediction challenges, enabling a broad assessment of the current state of nucleic acid structure prediction accuracy. The results confirm a continuing gap in accuracy between nucleic acid structure prediction and the near-experimental accuracy that is routinely achieved in protein structure prediction. Furthermore, automated methods remain behind the best human expert groups, despite the widespread exploration of automated deep learning methods. For RNA monomer modeling, which has been assessed in blind challenges like RNA-puzzles for over a decade, prediction accuracy has historically been heavily dependent on the quality of available 3D structure templates, and we observe no significant improvement in CASP16. Due to the observed template dependence of performance, future CASP16 assessments may consider the separation of assessment into template and free modeling, as was done historically for proteins^71,88^ prior to the advent of AlphaFold.

Nucleic acids are often “social” molecules, interacting with metabolites and proteins to transmit information. For the first time, CASP16 engaged the prediction community in new challenges in DNA structure prediction and RNA multimer prediction and assessed predictions of NA-protein and NA-ligand interfaces. For these targets, as with RNA monomer targets, prediction accuracy remains tied to the availability of 3D structure templates, and much progress is needed. The inclusion of nucleic acid targets in CASP’s model quality assessment category^89^ in the future, may benefit the community by encouraging more accurate self-assessment.

There are signs that evolutionary data improve nucleic acid structure prediction, as was the case for protein structure prediction starting even before the influx of deep learning^90^. Whether further advances in modeling RNA molecules, DNA molecules, and their complexes can be driven by more sophisticated use of evolutionary data, deep learning advances, or other emerging sources of high-throughput experimental data remains to be seen. A jump in accuracy may lie on the horizon, but it will apparently require not just importing strategies from protein modeling but developing novel approaches unique to nucleic acids.

## Data availability

The predicted models can be found at: https://predictioncenter.org/download_area/CASP16/predictions/RNA/; https://predictioncenter.org/download_area/CASP16/predictions/oligo/; https://predictioncenter.org/download_area/CASP16/predictions/hybrid/.

The group abstract and presentations can be found at: https://predictioncenter.org/casp16/doc/CASP16_Abstracts.pdf; https://predictioncenter.org/casp16/doc/presentations/Day-3/.

The data that support the findings of this study and code to replicate these results are openly available in a GitHub repository https://github.com/DasLab/CASP16_NA and at the CASP website, where the scores can be found at: https://predictioncenter.org/casp16/results.cgi?tr_type=rna; https://predictioncenter.org/casp16/results.cgi?tr_type=rna_multi; https://predictioncenter.org/casp16/results.cgi?tr_type=hybrid.

## Author Contributions

R.C.K., A.K., and R.D. conceptualized and designed the study. A.K. organized the target and prediction submission and calculated metrics for monomeric and multimeric targets. S.H., T.K., and R.C.K. curated the secondary structure algorithm predictions. S.H., R.D., and R.C.K. qualitatively reviewed models and conducted the base pair analysis. A.M.H. and R.C.K. conducted the motif analysis and MSA analysis. J.Z., R.Y. and Q.C. curated NA-protein hybrid targets and provided analysis methodologies for those targets. R.D. identified best available templates and structure pairs solved by independent groups. R.C.K. prepared the manuscript with input from all authors.

## Acknowledgments

We thank Gabriel Studer for sharing OpenStructure code updates and insights in early stages of assessment, Jérôme Eberhardt and Xavier Robin for sharing insights on ligand scoring with OpenStructure, Stanford Research Computing for expert administration of computing resources, and experimentalists providing nucleic acid structures as CASP16 targets. This work was supported by the National Institute for Health (NIGMS R01GM100482 to A.K., R35 GM122579 to R.D., R01 AI165433 to S.H., NIAID K99AI180984-01A1 to J.Z.), Stanford School of Medicine Dean’s Postdoctoral Fellowship (to A.M.H.), Stanford Bio-X (Bowes Graduate Student Fellowship to R.C.K.), Howard Hughes Medical Institute (HHMI) (to R.D.), the National Science Foundation (Grant No. 2330652 to R.D.), Endowed Scholars Program in UTSW (to Q.C.), and Welch Foundation (to I-2095-20220331 Q.C.). This article is subject to HHMI’s Open Access to Publications policy. HHMI lab heads have previously granted a nonexclusive CC BY 4.0 license to the public and a sublicensable license to HHMI in their research articles. Pursuant to those licenses, the author-accepted manuscript of this article can be made freely available under a CC BY 4.0 license immediately upon publication.

