## Supplemental Figures 1-9 for "Assessment of Nucleic Acid Structure Prediction in CASP16"


| | |
|---|---|
| Rachael C. Kretsch[1] | https://orcid.org/0000-0002-6935-518X |
| Alissa M. Hummer[2,3,*] | https://orcid.org/0000-0002-3023-2588 |
| Shujun He[2,4,*] | https://orcid.org/0000-0003-1010-536X |
| Rongqing Yuan[5,6,7] | https://orcid.org/0000-0001-5917-4505 |
| Jing Zhang[5,6] | https://orcid.org/0000-0003-4190-3065 |
| Thomas Karagianes[8] | |
| Qian Cong[5,6,9] | https://orcid.org/0000-0002-8909-0414 |
| Andriy Kryshtafovych[10] | https://orcid.org/0000-0001-5066-7178 |
| Rhiju Das[1,2,3,‡] | https://orcid.org/0000-0001-7497-0972 |

[1] Biophysics Program, Stanford University School of Medicine, Stanford, CA, USA.
[2] Howard Hughes Medical Institute, Stanford University, Stanford, CA, USA.
[3] Department of Biochemistry, Stanford University School of Medicine, Stanford, CA, USA.
[4] Department of Chemical Engineering, Texas A&M University, TX, USA.
[5] Eugene McDermott Center for Human Growth and Development, University of Texas Southwestern Medical Center, Dallas, TX, USA.
[6] Department of Biophysics, University of Texas Southwestern Medical Center, Dallas, TX, USA.
[7] Lyda Hill Department of Bioinformatics, University of Texas Southwestern Medical Center, Dallas, TX, USA.
[8] Eterna Massive Open Laboratory, Stanford, CA, USA.
[9] Harold C. Simmons Comprehensive Cancer Center, University of Texas Southwestern Medical Center, Dallas, TX, USA.
[10] Genome Center, University of California, Davis, CA, USA.

[*] Contributed equally

### Supplemental Tables

Each table below can be found in a tab in the supplied supplemental spreadsheet.

**Supplemental Table 1: List of unscored models.**

**Supplemental Table 2: Model scores.**

**Supplemental Table 3: The best prediction and template for RNA blind prediction challenges.**

**Supplemental Table 4: Group sum Z-scores for 3D structures and mean F1 scores for base pairs and tertiary motifs.**

**Supplemental Table 5: Pairs of independently determined structures of the same RNA sequence.**

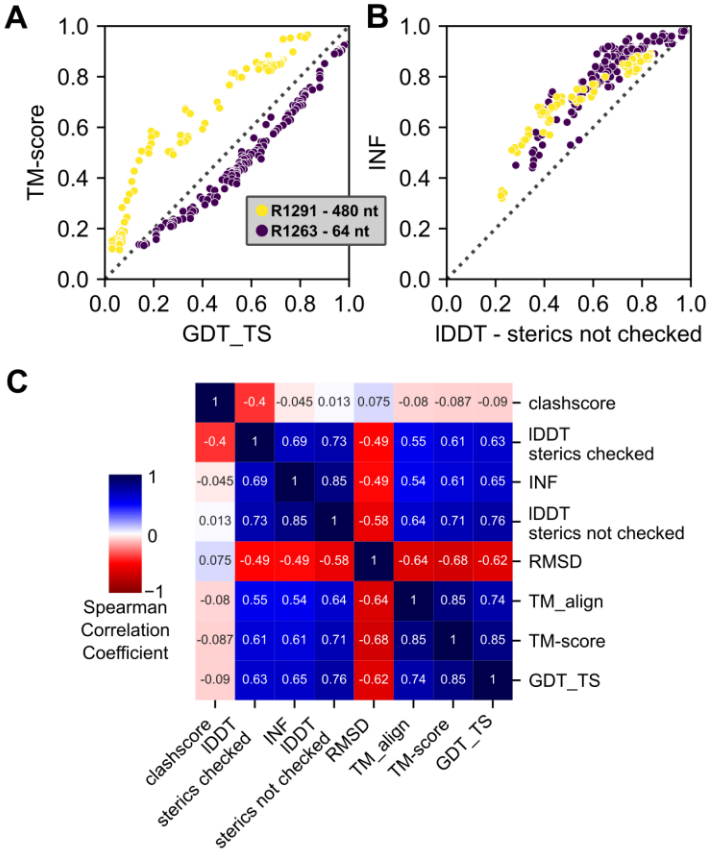


**Supplemental Figure 1: Analysis and comparison of scoring metrics.** A representative short target, R1263 (64 nt), and long target, R1291 (480 nt) are chosen to illustrate the length-dependence of metrics. (**A**) The TM-score for all predicted models is plotted against the GDT_TS, showcasing the different behaviour of the two-scores at different sequence lengths. (**B**) The INF for all predicted models is plotted against the lDDT without the steric check, showing high correlation and similar trends across the length scale. (**C**) The Spearman correlation coefficient calculated for all pairs of metrics for every target separately and then averaged. The Spearman correlation coefficient is labeled and colored from strong negative correlation, red, to strong positive correlation, blue. The scores are clustered by the Euclidean distance; the absolute value of the correlations is taken before calculating the distance to omit the directionality of scores (e.g. a lower clashscore, but a higher lDDT is better).

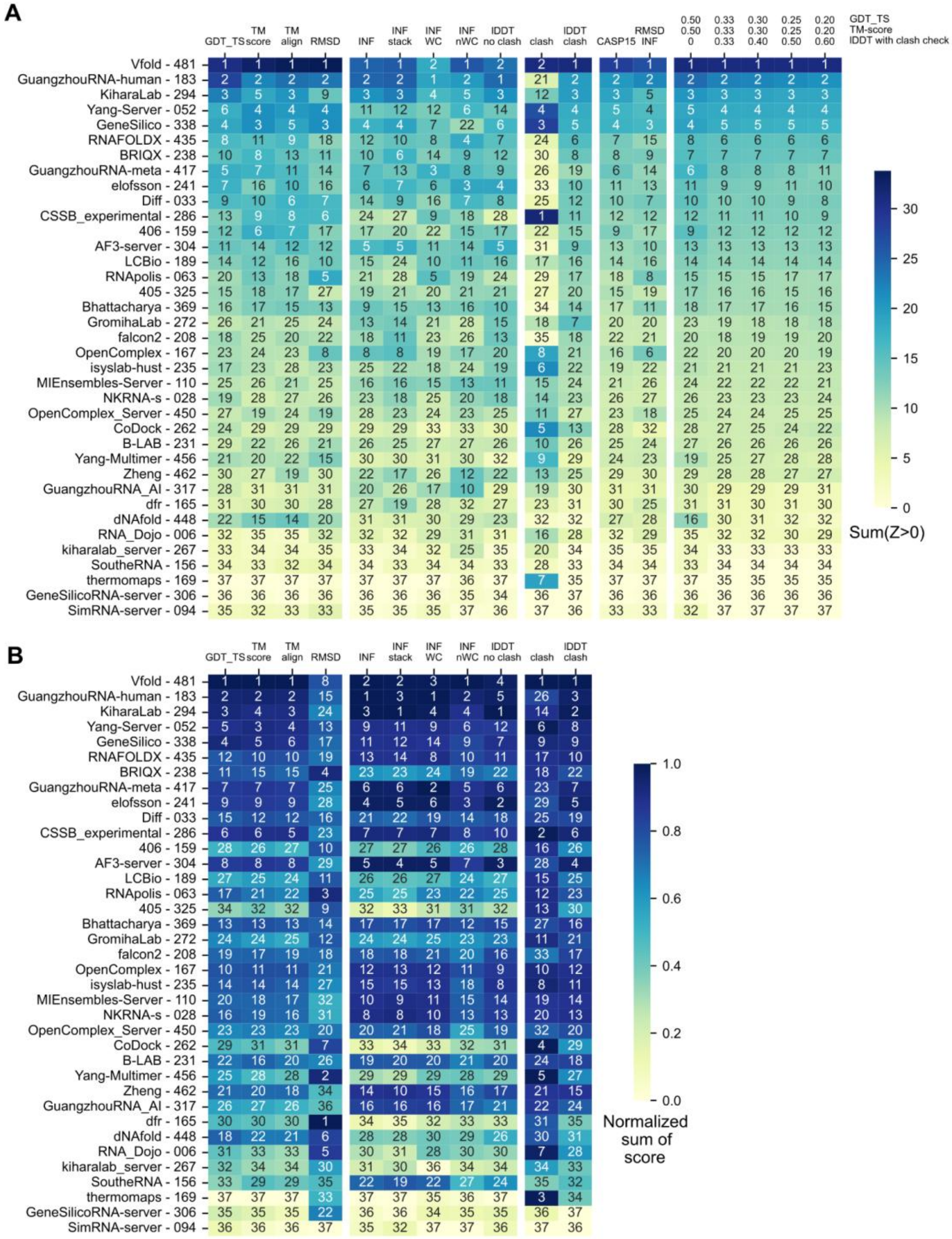


**Supplemental Figure 2: Robustness of ranking to the scoring used.** The groups were limited to those that submitted at least one model for >60% of the targets. (**A**) Groups are ordered by their ranking in CASP16. For each of the scoring methods (columns) the rank of the group is annotated and the Sum(Z>0) is represented by the color of the box. All scores in (A) are Sum(Z>0) where a Z-score is calculated for each target and then all summed together, not penalizing for negative Z-scores. The leftmost columns are global fold metrics, the next groups of columns are local environment scores, then the third group are metrics that consider atomic clashes. The CASP15 score is $\frac{1}{3} Z_{TM\text{-}score} + \frac{1}{3} Z_{GDT\_TS} + \frac{1}{8} Z_{INF} + \frac{1}{8} Z_{lDDT} + \frac{1}{12} Z_{clash}$ and the RMSF-INF column is calculated by evenly weighted RMSD and INF Z-scores which RNA puzzles use. The rightmost group of columns displays the scoring with different weights of our chosen metrics; CASP16 scoring is the center column. (**B**) The same metrics are plotted for each group, however, instead of Z-scores, the raw scores are summed and the summed scores are used to rank the groups. For display purposes, these summed scores are normalized to between 0 (worst) and 1 (best).

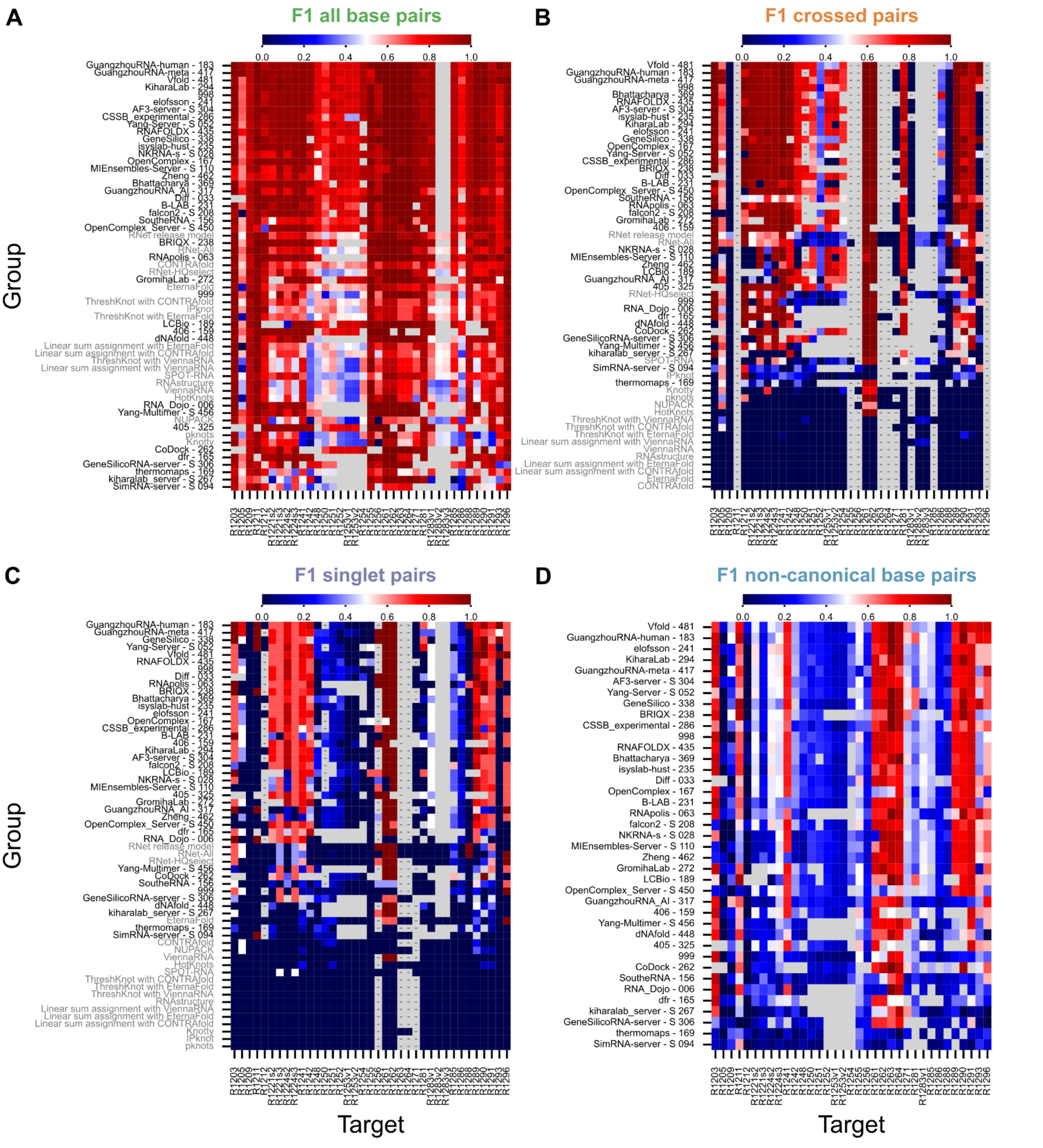


**Supplemental Figure 3: Base-pairing prediction accuracy.** For all groups and select secondary structure prediction algorithms (labeled gray), (**A**) the set of all base pairs, (**B**) crossed base pairs, (**C**) singlet base pairs, and (**D**) non-canonical base pairs were compared to the target. The F1-score is plotted for the best prediction for each target from each group or secondary structure prediction algorithm. The predictors are ordered by average F1-score. Groups that did not participate in a target would receive a score of 0 for that target. Gray boxes indicate a group did not participate in that target and a gray box with a dash indicates the group did participate and correctly identified that interaction was not present in the target.

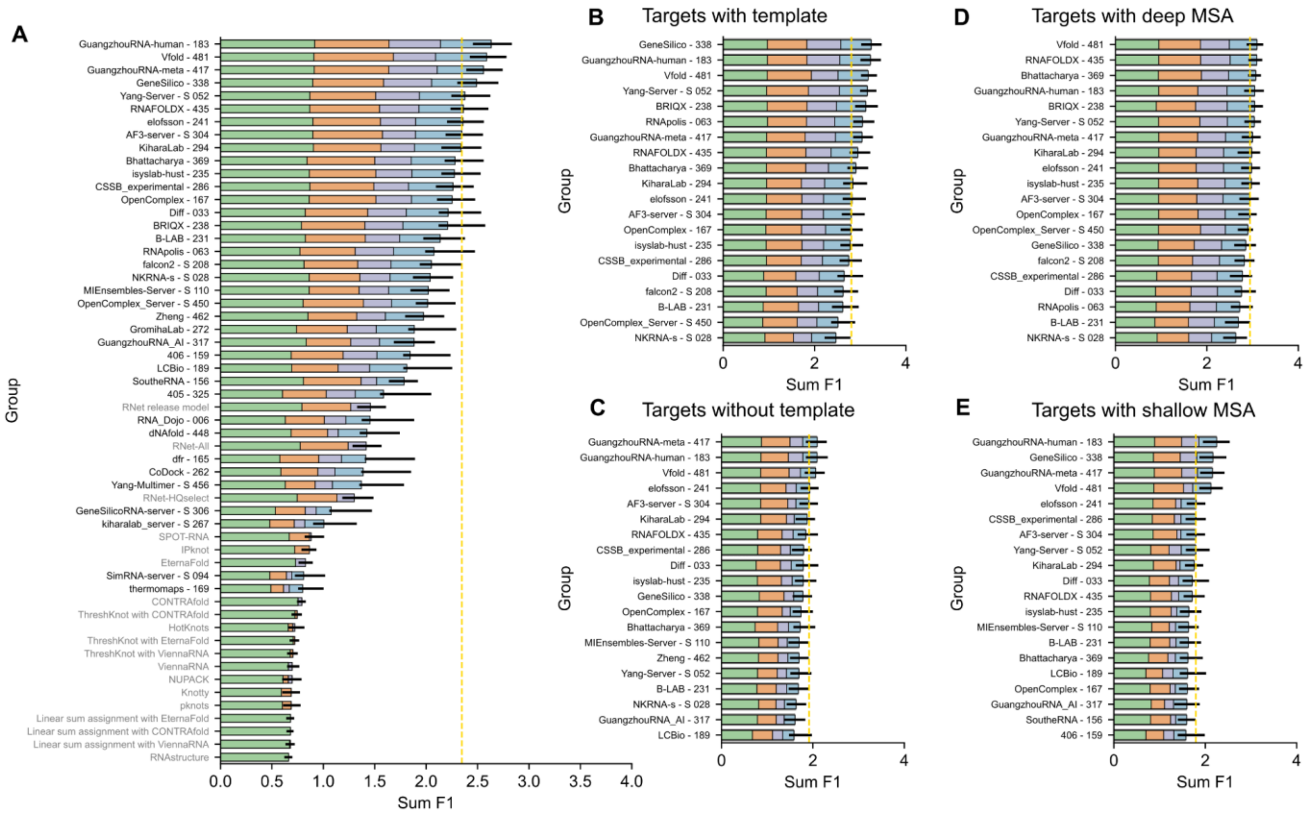


**Supplemental Figure 4: Ranking of groups by base-pairing prediction accuracy.** For all groups and select secondary structure prediction algorithms (labeled gray), (**A**) the ranking of groups is taken after summing across the four F1-scores in **Supplemental Figure 3**. The average F1-score of all base pairs (green), crossed base pairs (orange), singlet base pairs (purple), and non-canonical pairs (blue) are plotted. There are a total of 2,880 canonical base-pairs, 147 canonical singlet base-pairs, 935 crossed canonical base-pairs, and 1,439 non-canonical base-pairs in all targets. Note F1 also penalizes over-prediction, so the number of base-pairs predicted also affects F1. The targets are split to show ranking for (**B**) only targets with a template (TM-align > 0.45), (**C**) targets without a template, (**D**) targets with deep MSAs (Neff > 130) and (**E**) targets with shallow MSAs. The performance of AlphaFold 3 Server is shown with a gold dotted line. Error bars represent the 68.2% confidence interval.

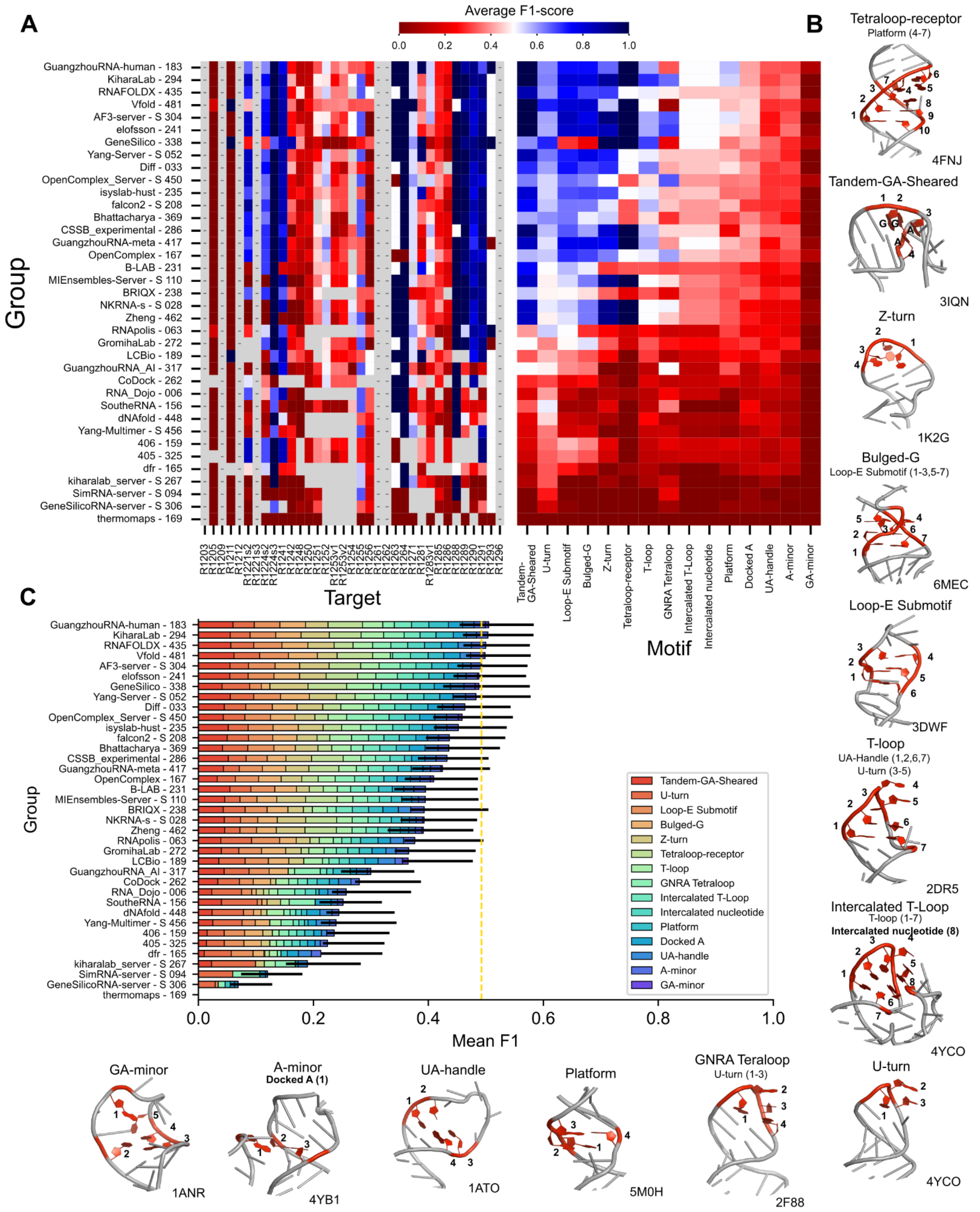


**Supplemental Figure 5: RNA motif prediction accuracy.** For all predictions, the set of nucleotides involved in various RNA motifs are calculated and compared to the target. (**A**) The average F1-score is calculated for the best score for each group. On the left, the F1-score is average across all motif-types. On the right, the F1-score is average across all targets. (**B**) Examples of motifs, labeled by the order in the rna_motif reports them. (**C**) RNA motif prediction results are summarized. The average F1-score across all targets. The bar is split into contributions from each motif type. The performance of AlphaFold 3 Server is shown with a gold dotted line. Error bars represent the 68.2% confidence interval.

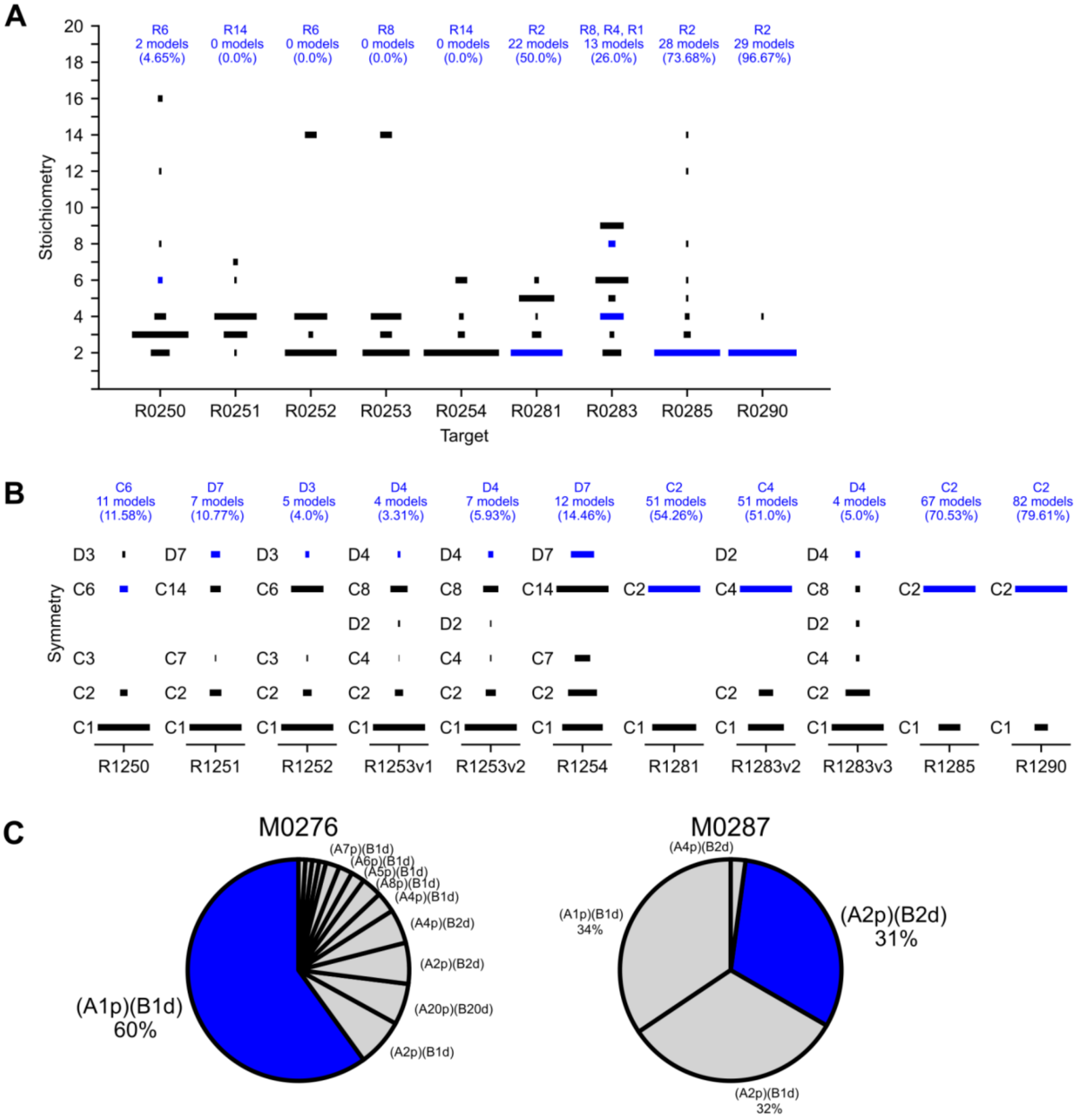


**Supplemental Figure 6: Accuracy of stoichiometry and symmetry predictions.** (**A**) The stoichiometry of predictions for round 0 RNA-homoligomer targets are represented by bars proportional to the number of predictions submitted with each stoichiometry. The stoichiometry of the target is labeled in blue. (**B**) The symmetry of predictions for round 1 RNA-homoligomer targets are represented by bars proportional to the number of predictions submitted with each stoichiometry. The symmetry is labeled on the left of each plot, with the top representing the largest dihedral symmetry when applicable, the second representing the largest cyclical symmetry, the bottom representing no symmetry and all other symmetries in the middle. The symmetry of the target is labeled in blue. (**C**) The predicted stoichiometry round 0 hybrid targets are represented in a pie chart where the size of each slice is proportional to the number of predictions submitted with each stoichiometry. The first number represents the number of protein chains predicted, the second number represents the number of DNA chains predicted. The stoichiometry of the target is labeled in blue. Models that did not predict any DNA are removed.

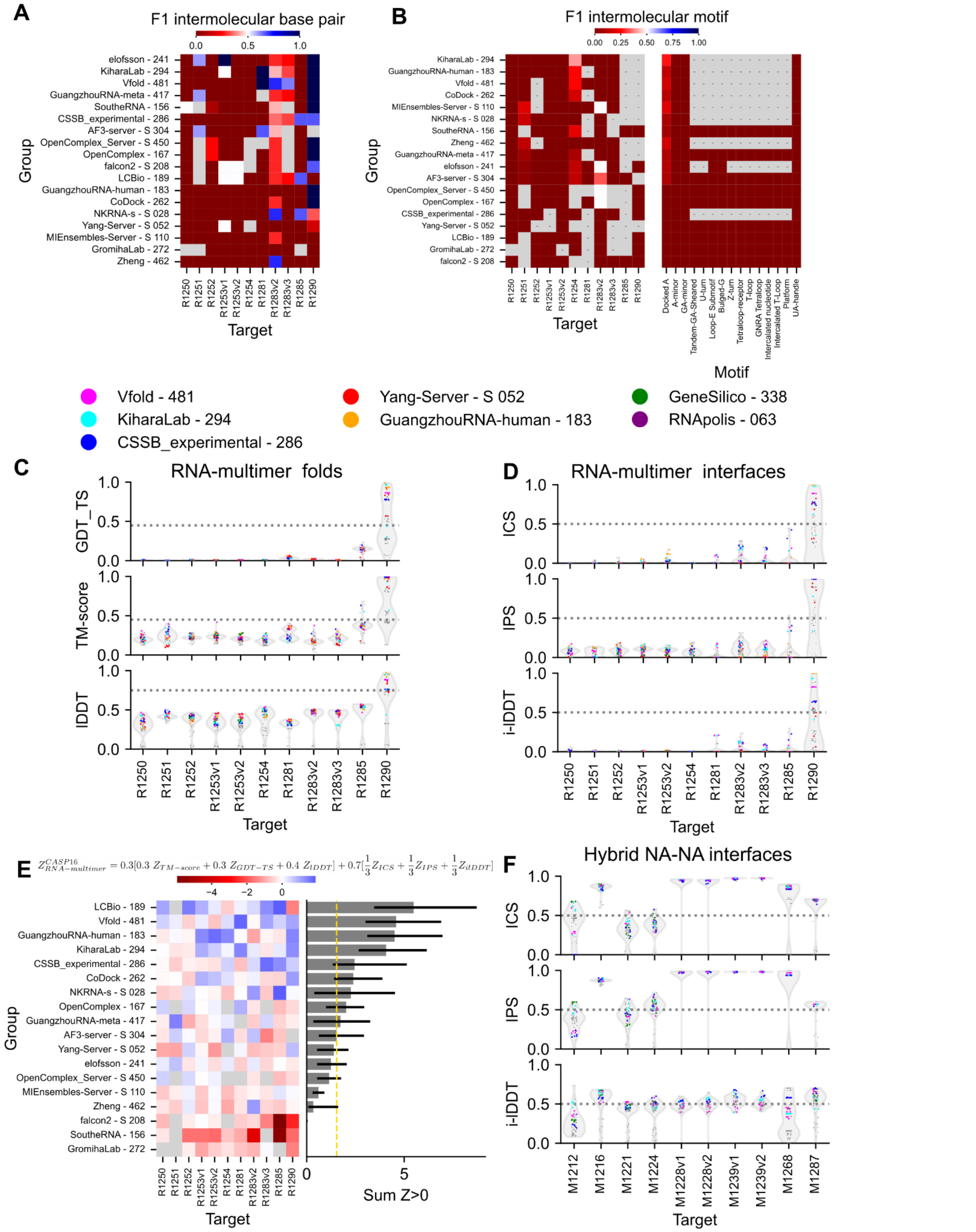


**Supplemental Figure 7: Accuracy of NA-NA interface and RNA multimer prediction.** (**A-B**) The F1-score of intermolecular base pairs as shown in **Supplemental Figure 3A** and motifs as shown in **Supplemental Figure 5A** are displayed. (**C**) The scores measuring the accuracy of the fold of the entire complex for RNA multimer targets is shown The distribution of scores is displayed in the violin plot, with individual scores displayed with gray dots, colored for select teams. (**D**) The interface scores for NA-NA interfaces for RNA multimer targets are shown in the same style as (C). (**E**) The participating groups are ranked according to $Z^{CASP16}_{RNA-multimer}$ . The sum of non-negative Z-score over all targets is displayed right, with 68.2% confidence interval as the error bar. (**F**) The interface scores for NA-NA interfaces for hybrid targets are shown in the same style of (C).

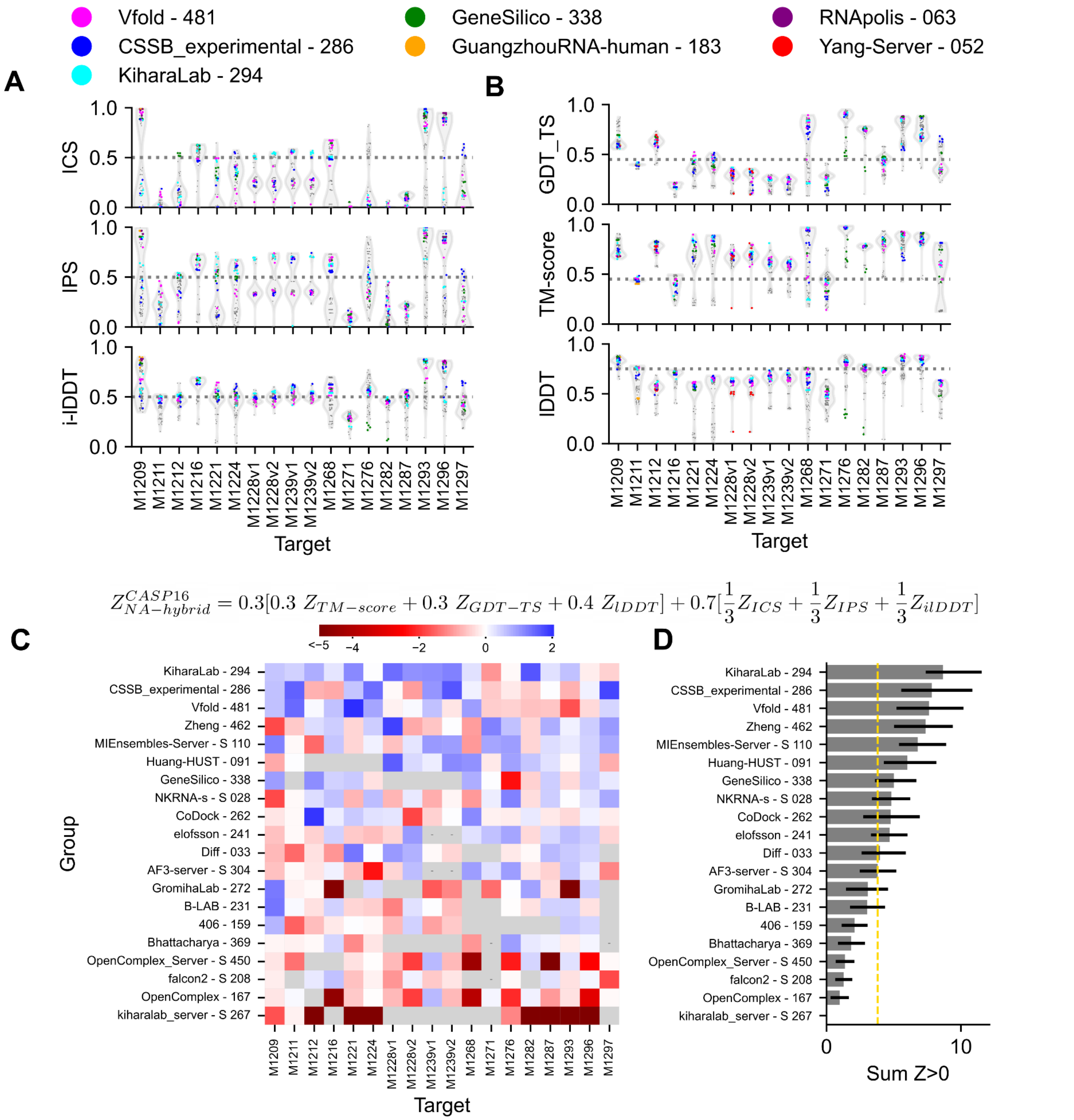


**Supplemental Figure 8: Accuracy of NA-protein interface and hybrid complex prediction.** (**A-B**) The interface scores for NA-protein interfaces are shown for hybrid targets. The distribution of scores is displayed in the violin plot, with individual scores displayed with gray dots, colored for select teams. (**B**) The scores measuring the accuracy of the fold of the entire complex for hybrid targets is shown in the same style as (A). (**C**) The participating groups are ranked according to $Z^{CASP16}_{NA-hybrid}$ . (**D**) The sum of non-negative Z-score over all targets is displayed right, with 68.2% confidence interval as the error bar and AlphaFold 3 Server performance marker with gold dotted line.

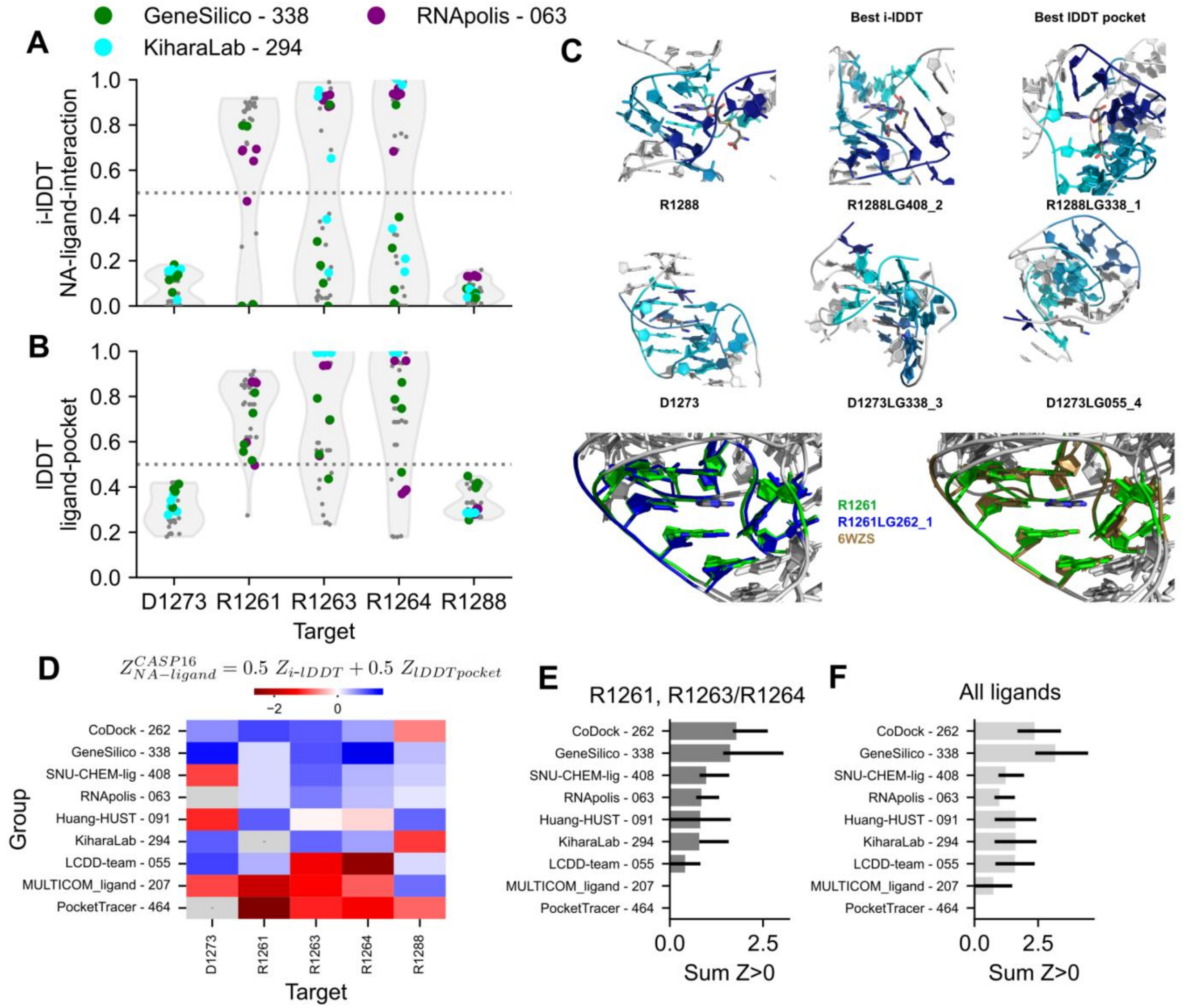


**Supplemental Figure 9: Ligand prediction results.** (**A**) The i-lDDT of the NA-ligand interface and (**B**) lDDT of the nucleotides that make up the pocket are plotted for all 5 targets. A violin plot displays the spread of scores. Individual predictions are displayed with scatter points and key groups are colored. (**C**) Top scoring predictions are displayed. For D1273 (DNA aptamer with dopamine) and R1288 (SAMURI ribozyme with S-adenosyl methionine cofactor), the target and predicted ligand are aligned and the nucleotides interacting (< 6 Å) and the ligand in the target are colored blue. The same residues are colored in the predictions. No group predicted the ligand pocket and NA-ligand interactions like stacking, are generally under-predicted. For R1261, representative of the ZTP-riboswitch targets, the best predicted ligand pocket (blue, left) and the best template pocket (brown, right) are aligned to the target (green). Only the part of the ligand which makes the majority of NA-ligand interaction is displayed for clarity. Predictions were high quality, as expected from template-based modeling. (**D**) The $Z^{CASP16}_{NA-ligand}$ is plotted for every group and target. (**E-F**) For every group, the sum of Z-scores that are greater than 0 is summed, and the 68.2% confidence interval is displayed. When the pocket for the ligand is very poorly formed, as in all predictions for R1288 and D1273, it can be misleading and difficult to interpret accuracy of ligand pose, hence the final ranking excludes these targets. AlphaFold 3 did not participate in this category due to the limited ligand selection of the server, so the baseline is not displayed. No servers participated in this category so no groups are labeled with an “S”.
